# Single-cell genetic heterogeneity linked to immune infiltration in glioblastoma

**DOI:** 10.1101/2021.09.20.461080

**Authors:** Kacper A. Walentynowicz, Dalit Engelhardt, Shreya Yadav, Ugoma Onubogu, Roberto Salatino, Cristina Vincentelli, Thomas O. McDonald, Franziska Michor, Michalina Janiszewska

## Abstract

Glioblastoma (GBM) is the most aggressive brain tumor with a median survival of ~15 months. Targeted approaches have not been successful in this tumor type due to the large extent of intratumor heterogeneity. Mosaic amplification of oncogenes suggests that multiple genetically distinct clones are present in each tumor. To uncover the relationships between genetically diverse subpopulations of GBM cells and their native tumor microenvironment, we employed highly multiplexed spatial protein profiling, coupled with single-cell spatial mapping of fluorescence in situ hybridization (FISH) for *EGFR, CDK4*, and *PDGFRA*. Single-cell FISH analysis of a total of 35,843 single nuclei (~2,100 per tumor) revealed that tumors in which amplifications of *EGFR* and *CDK4* more frequently co-occur in the same cell exhibit higher infiltration of CD163^+^ immunosuppressive macrophages. Our results suggest that high throughput assessment of genomic alterations at the single cell level could provide a measure for predicting the immune state of GBM.

## INTRODUCTION

Glioblastoma multiforme (GBM) is the most heterogeneous and aggressive brain malignancy, with an average survival of 15-18 months post-diagnosis (Wen et al., 2020). Even through GBM’s frequency of 3 cases per 100,000 people in the U.S is moderate, this disease causes high morbidity and mortality, as only 6.8% patients survive beyond 5 years post-diagnosis (Wen et al., 2020). This poor prognosis has not changed significantly in recent decades. Combination of chemotherapy and radiation is the only treatment providing significant clinical benefit, yet on average it prolongs patients’ life by only 4 months (Wen et al., 2017). Despite a growing understanding of the disease biology, therapies targeting molecular features in GBM have been failing in clinical trials (Aldape et al., 2019).

Challenges to effective treatment of GBM are fueled by the large extent of its intratumor heterogeneity. Recent advances in molecular profiling have unraveled the complexity underlying GBM cellular diversity at the genetic, epigenetic and phenotypic levels. Genetically, GBMs are characterized by a complex mutational landscape with a large degree of inter- and intratumor heterogeneity (J.-K. Lee et al., 2017; McKenna et al., 2013; Sottoriva et al., 2013). Gains in chromosome 7 and loss of chromosome 10 are common events predicted to arise early in tumor evolution (Ozawa et al., 2014). Frequent amplifications of several receptor tyrosine kinases (RTK), including EGFR, PDGFRA and MET, have been explored as potential therapeutic targets in GBM (Wen et al., 2020). However, despite successful trials in other malignancies, tyrosine kinase inhibitors have not been found to provide significant benefit to GBM patients (Rich et al., 2004; Wen et al., 2020; 2006). In addition to the adaptability of cellular signaling in response to RTK inhibition, genetic intratumor heterogeneity further contributes to GBM resistance (Nicholson and Fine, 2021). In particular, RTK-encoding gene amplifications, *PDGFRA, EGFR* and *MET*, have been shown to frequently co-occur in the same GBM sample, yet not in the same cells (Snuderl et al., 2011; Szerlip et al., 2012). Thus, genetic mosaicism of GBM is one of the key obstacles for effective treatment of these tumors.

Inter- and intratumor heterogeneity of gene amplifications is also linked to the diversity of phenotypes identified in GBM. Initial transcriptional profiling of bulk tumor tissue revealed that GBMs can be classified into three subtypes: classical-like (CL), proneural (PN) and mesenchymal (MES) (Verhaak et al., 2010; Q. Wang et al., 2017). Each of these transcriptional subtypes is associated with increased frequency of specific genetic alterations. *EGFR* amplification and loss of *CDKN2A/B* are typical for classical-like GBM, *CDK4* and *PDGFRA* amplifications for proneural GBM, and *NF1* loss is associated with a mesenchymal transcriptional program (Q. Wang et al., 2017). Despite this discovery, the restriction to the proneural tumor type in clinical trials for PDGFRA inhibition did not increase the treatment success rate (Wen et al., 2020). Advances in single cell transcriptomics revealed that each GBM tumor, regardless of subtype reported by bulk profiling, is in fact a mixture of cells belonging to different subtypes (Garofano et al., 2021; Neftel et al., 2019; Patel et al., 2014). One of the most recent classifications of cells based on single cell expression profiling identified four major cellular states (Neftel et al., 2019). This classification also points to gene amplifications as potent drivers of intratumor heterogeneity, as each of the cellular states is associated with a distinct genetic alteration: oligodendrocytic precursor cell-like (OPC-like) with *PDGFRA* amplification, neural progenitor cell-like (NPC-like) with *CDK4* amplification, astrocytic cell-like (AC-like) with *EGFR* amplification and mesenchymal-like (MES-like) with NF1 loss (Neftel et al., 2019). While transitions between these cell types can occur, each genetic driver favors a particular transcriptional state of the cell. Thus, assessment of copy number alterations (CNA) for *EGFR, CDK4* and *PDGFRA* amplifications at the single cell level *in situ* may provide information about cellular diversity within distinct tumor microenvironments.

*In situ* heterogeneity of GBM, observed as a large extent of variability of histological features, is characteristic for this tumor type and forms the root of the name “multiforme”. The disordered tissue structure as well as non-uniform contrast enhancement in magnetic resonance imaging (MRI) point to a high degree of macroscopic diversity within these tumors (O’Connor et al., 2015). Characteristic pathological features of GBM include areas of pseudopalisading, organized alignment of viable cells surrounding the necrotic regions, and microvascular proliferation (Tomaszewski et al., 2019). These different tumor microenvironmental types are often intermixed in the tumor tissue. While the mechanisms that generate these features are not fully understood, it is thought that pseudopalisades are created by tumor cells rapidly migrating away from a necrotic site (Brat et al., 2004). These necroses arise due to vaso-occlusion caused by intravascular thrombosis and cells in pseudopalisades activate the HIF1alpha hypoxia response (Rong et al., 2006; Tehrani et al., 2008). Interestingly, the perinecrotic areas and perivascular regions, constituting drastically different environments, were both shown to support GBM stem-like cells, rare subpopulations with increased epigenetic plasticity and potentially higher resistance to standard treatment (Calabrese et al., 2007; Janiszewska et al., 2012; Lathia et al., 2015; Leder et al., 2014; Prager et al., 2019). Constant changes in the tumor microenvironment between a highly oxygenated, nutrient-rich state and a hypoxic state with collapsing vasculature could significantly affect the survival of distinct cell subpopulations in these microenvironments. Thus, to survive, GBM cells may need to adapt to these new selective pressures. Yet, this adaptation is not necessarily cell-autonomous; instead, the maintenance of intratumor heterogeneity in GBM may be dependent on interactions between genetically diverse GBM cell subpopulations and may be driven by differences in and interactions with specific microenvironments.

Since the four major subtypes of GBM cells, while remaining plastic, are largely determined by underlying genetic aberrations, we aimed to study how genetically constricted populations interact with each other in the context of the tumor microenvironment of intact clinical tissue specimens. To this end, we performed a spatial analysis of protein markers and single cell quantification of *CDK4, EGFR* and *PDGFRA* copy number in 17 GBM samples in multiple areas of each biopsy. The increased power of single cell automated quantification allowed us to identify tumors with high odds ratio of *CDK4* and *EGFR* co-amplification in the same cells. We found that the microenvironment of tumors with co-amplified cells was significantly enriched in infiltrating immune cells as compared to tumors with fewer co-amplified cells. Our results suggest that a quantification of genetic heterogeneity based on large-scale single cell measurements in individual biopsies could help identify biologically distinct GBM tumor groups.

## RESULTS

To investigate the relationship between the spatial localization of cells with different genotypes and their microenvironmental preferences, we constructed a tissue microarray (TMA) from 17 formalin-fixed paraffin-embedded (FFPE) GBM cases (**Supplemental Table 1**). From each tumor block, 3-4 cores were randomly selected (61 in total) to investigate the patterns of local intratumor heterogeneity within each tumor biopsy (**Fig. 1a**). Since these samples were of variable age and embedded in paraffin blocks, they could not be used for transcriptomic studies; however, formalin-fixed paraffin embedding and prolonged storage in ambient temperature does not impede immunohistochemistry or DNA-based studies. Thus, we used our multi-region TMA of intact tissue samples from each tumor for single cell assessment of select genomic features by fluorescence *in situ* hybridization (FISH) and specific-to-allele PCR-FISH (STAR-FISH) as well as protein expression using NanoString’s GeoMx Digital Spatial Profiling (DSP) platform.

**Figure 1.**
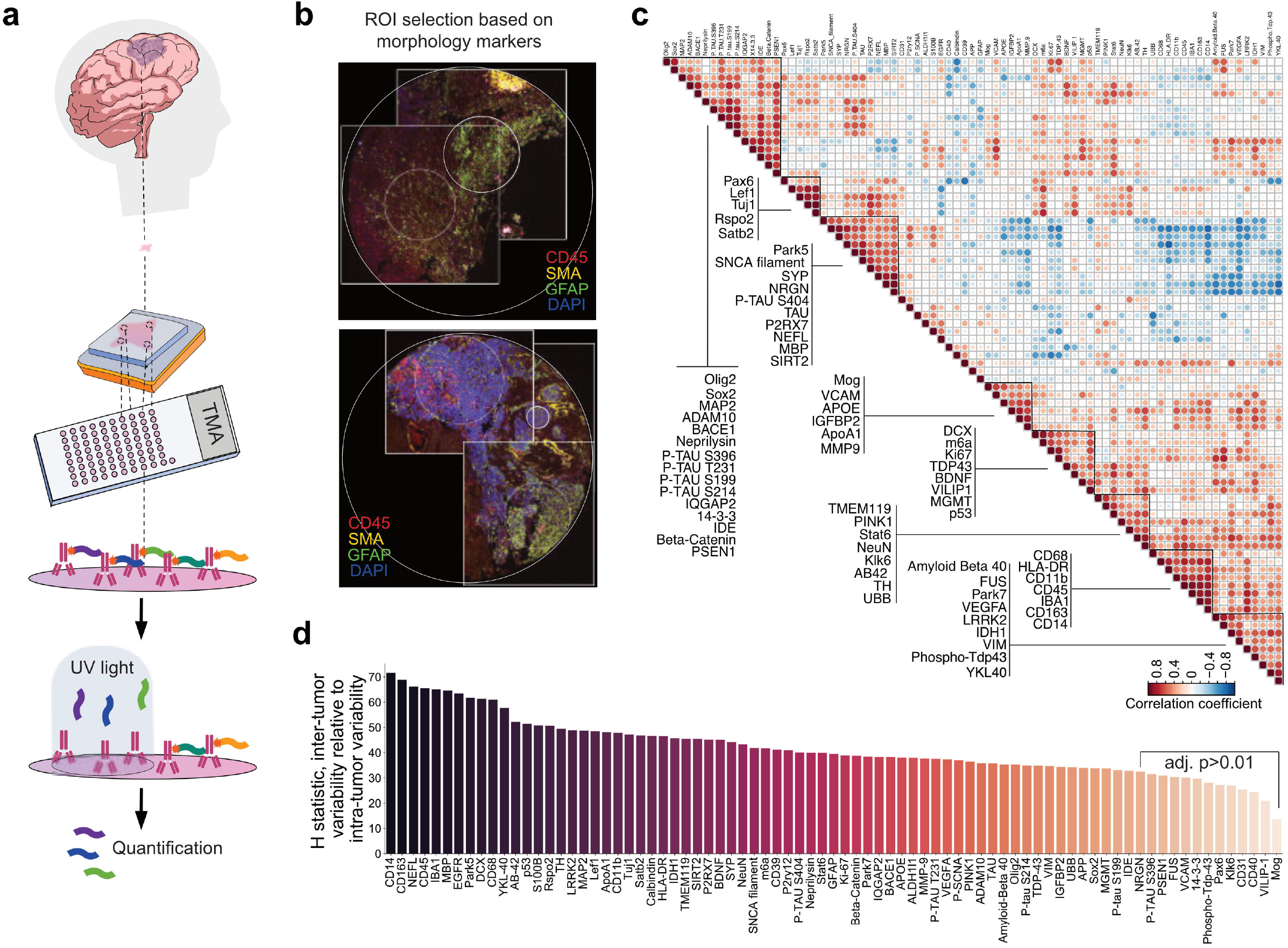
Spatial profiling reveals regional heterogeneity in protein expression between distinct areas of the same GBM biopsy. **a)** Schematic of tumor microarray (TMA) construction and digital spatial profiling (DSP) for protein marker expression. GBM biopsies from 20 patients were embedded in paraffin blocks. From each block 3-4 cores were punched to construct a multiregion TMA. In DSP assay, staining with 79 protein marker-specific antibodies labeled with photocleavable oligo-tags were used. UV light exposure of a region of interest (ROI) releases the tags, which are then quantified for each ROI separately. **b)** Example of two cores with low (left) and high (right) heterogenous staining used to select regions-of-interest for DSP. Large circle – full core size (1mm diameter), small circles – selected ROIs. **c)** Spearman correlations between all profiled markers. Groups of markers with the strongest correlations are marked. **d)** Relative inter-tumor to intra-tumor variability, as indicated by the H statistic (see **Methods**), in the expression of each protein.

### GeoMx DSP data reveals local heterogeneity in tumor microenvironmental marker expression

To characterize the extent of variation in the local GBM tumor microenvironment within distinct regions of the same tumor biopsy, we employed a spatial profiling platform based on highly multiplexed immunostaining with oligonucleotide-tagged antibodies (**Fig. 1a**). GeoMx DSP allows quantification of oligo-tags released by UV-illuminated regions of interest (ROI), which can be selected based on immunofluorescent staining for a few markers. DSP enables simultaneous antibody staining for tens of markers and control antibodies, providing more robust quantification compared to fluorescence-based methods. In our assay, multicolor immunofluorescent staining with markers of glial cells (GFAP), immune cells (CD45) and vasculature (α-SMA) was used to select ROIs within each TMA core (**Supplemental Fig. 1a**). Despite the small size of each core (1 mm in diameter), many of the cores display varied morphology and expression levels for GFAP, CD45 and α-SMA within the core (**Fig. 1b**). We selected 96 regions of interest (ROI) out of the 61 cores present on the TMA (see **Methods**), with multiple (2-3) ROIs in cores with heterogeneous immunofluorescent staining (**Supplemental Fig. 1a**). These ROIs were then subjected to GeoMx digital spatial profiling for 79 protein markers associated with glioblastoma biology, neurobiology and immunology (**Fig. 1a,b, Supplemental Table 2**). The data were normalized to positive controls from the External RNA Controls Consortium (ERCC) and IgG isotype controls per ROI to account for differences in hybridization efficiency, background signal, cellularity, and size of ROIs (see **Methods**). We then performed a multi-level analysis of correlation in expression patterns across tumors and a quantification of intra- versus inter-tumor expression heterogeneity based on ROI- and core-specific expression patterns.

We found that the expression levels of the immune markers CD68, HLA-DR, CD11b, CD45, IBA1, CD163, and CD14 were correlated across all tumors (mean Spearman correlation 0.76, range 0.57-0.92, for clusters; **Fig. 1c**). A similar trend was observed for the TAU protein and its phosphorylated versions, except for phosphor-TAU S404 (Spearman correlation >0.69; **Fig. 1c**). Negative correlation was observed between clusters of immune (including CD68, HLA-DR and CD11b) and neuronal markers (Park5, SNCA-filament, SYP, MBP, among others), with a cluster-wise Spearman coefficient of −0.48, p = 0.052 (see **Methods**). A strong correlation was also observed between the expression of Lef1 and Rspo2, two proteins of the Wnt signaling pathway, and Satb2, a protein of the TCF/LEF pathway (Spearman correlation coefficients 0.88 and 0.93, respectively, and a Spearman correlation coefficient of 0.74 between Lef1 and Rspo2). These proteins have been implicated in stemness and promotion of a cancer stem cell phenotype (Clevers et al., 2014; Huang et al., 2020; Rheinbay et al., 2013).

In several cases, expression levels of the 79 markers were variable between cores derived from the same tumor in several cases (**Supplemental Fig. 1b**). Since the TMA cores represent spatially distant regions of the same tumor fragment with possibly different compositions, we assessed the variance of protein expression between ROIs from distinct tumors relative to the variance within tumors using the Kruskal-Wallis H test. Overall, we found that the variance of our 79-protein markers expression panel between tumors exceeded the variance within tumors (**Fig. 1d**). Particularly high inter-tumor relative to intra-tumor variance was observed for several immune related proteins (CD45, HLA-DR, IBA1, P2RX7, YKL-40), as well as for EGFR and p53, which are both frequently overexpressed and mutated in human GBM (Barthel et al., 2019; McKenna et al., 2013) and can drive tumorigenesis in animal models of glioma (J. H. Lee et al., 2018; Z. Wang et al., 2020). The neuronal markers NEFL, MBP, and Park5 also exhibited higher relative inter-tumor variance. On the other hand, vascular cell adhesion molecule (VCAM), and CD31, both associated with endothelial cell function and tumor vascularization (Weis and Cheresh, 2011), exhibited comparatively high expression variance between regions within the same tumor sample, suggesting that within a single biopsy of GBM there is high variability in blood vessel distribution (**Supplemental Fig. 2a,b**) and even a single biopsy is likely to contain differentially vascularized niches.

This result prompted us to assess the histopathological features of each TMA core to identify which protein markers may be specific for different tissue compositions (**Supplemental Fig. 2)**. Tissue composition assessment based on hematoxylin/eosin staining confirmed expected variation in hemorrhage and necrosis areas within cores from the same tumor (**Supplemental Fig. 2c**). However, classification of each core according to the most dominant histopathological feature did not reveal protein expression correlation patterns (**Supplementary Fig. 2d**), possibly as a consequence of cores containing mixed features.

### Characterization of single cell genetic heterogeneity in GBM multicore TMA

Amplifications of *EGFR, PDGFRA*, and *CDK4* occur frequently in GBM and are often present in the same tumor in different cellular populations. Each of these genetic alterations is associated with driving a particular transcriptional state (Neftel et al., 2019). The analysis of the spatial localization of cells harboring these genetic drivers could therefore be revealing of their microenvironmental preferences and interactions. To address this question, we performed multiplexed FISH for *EGFR, PDGFRA* and *CDK4* on our multi-region TMA (**Fig. 2a, Supplemental Fig. 3a**). Using an automated single cell FISH signal counting platform, we quantified the number of FISH signals in each individual nucleus imaged (n=35,843 nuclei, on average 2,335 nuclei per tumor, 216 nuclei per image, 166 images). Amplification of a gene was called when ≥ 6 FISH signals of a respective probe were found in a nucleus (see **Methods**). We found that *PDGFRA* amplification was relatively rare, detected in less than 2% of all cells analyzed, and we therefore focused on the amplification patterns of *EGFR* and *CDK4*, independently of *PDGFRA* amplification status. Each cell was assigned to one of the following genotypes: E (*EGFR* amplified and not *CDK4* amplified), C (*CDK4* amplified and not *EGFR* amplified), EC (*EGFR* and *CDK4* amplified), and N/O (no amplification of either *EGFR* or *CDK4*). We observed significant variation in the overall frequency of cells of each amplification genotype within each TMA tumor (**Fig. 2b-d**). Although several tumors had relatively high E cell frequency, C and EC cells were found in most of the analyzed samples. A calculation of the Shannon index of diversity (Magurran, 2005) of genotype prevalence within each image (**Fig. 2d**) shows variation between distinct cores taken from the same biopsy, further suggesting that the extent of heterogeneity differs within relatively small tumor regions.

**Figure 2.**
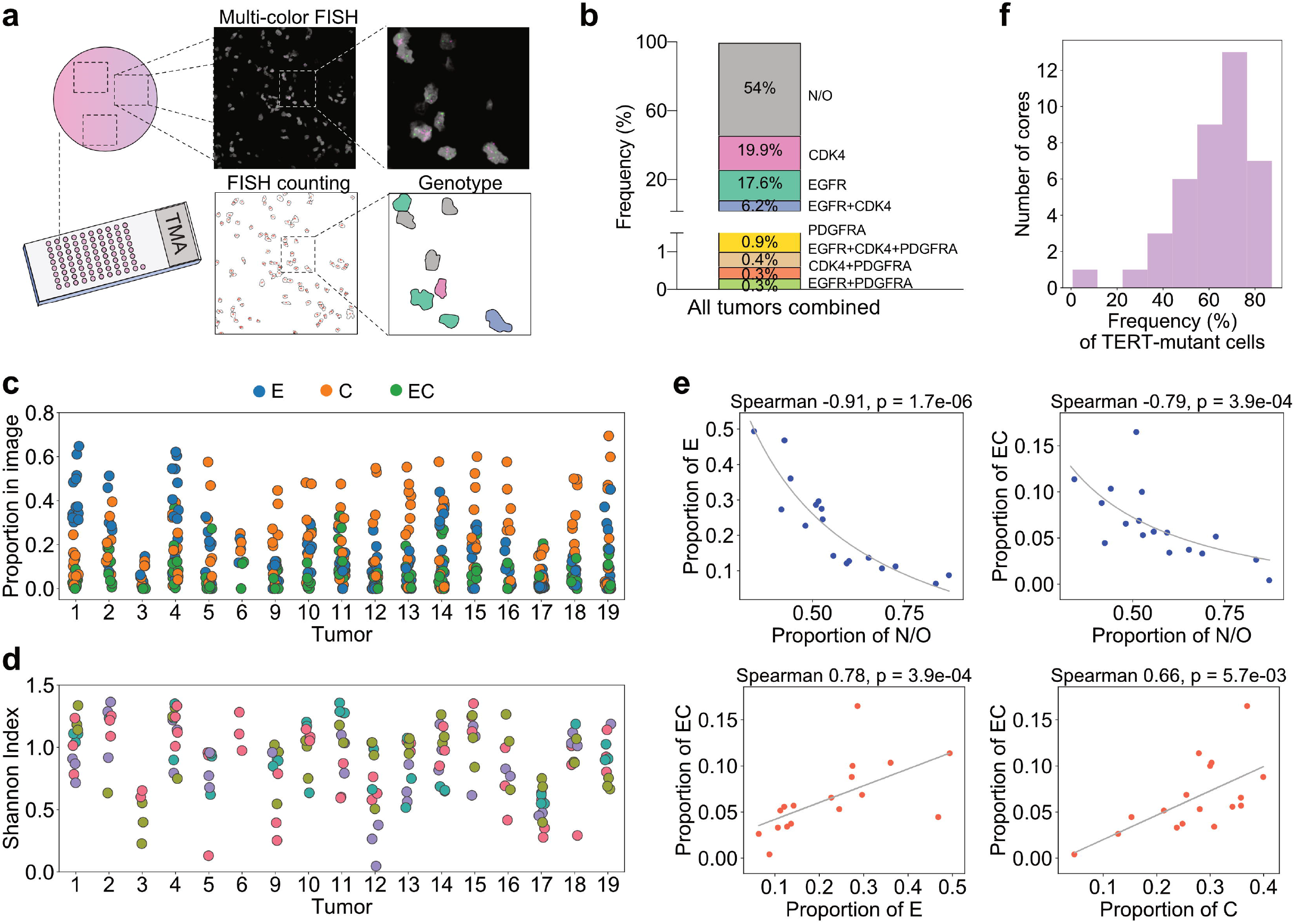
Single-cell characterization of EGFR, PDGFRA, CDK4 amplifications and hTERT promoter mutations in a multiregion GBM TMA. **a)** Summary of the workflow for FISH signal quantification in individual nuclei. Three images were taken per each TMA core and each imaged area was segmented, based on the nuclear outlies for counting of each FISH signal in individual nuclei. Based on presence of gene amplifications, each nucleus is classified as a distinct genotype. **b)** Overall frequency of cells with FISH-based genotypes. **c)** Proportion of cells with the E, C, and EC genotypes shown in individual images in each tumor. E – cells with amplified *EGFR* and no *CDK4* amplification, C – cells with amplified *CDK4* and no *EGFR* amplification, EC – cells with amplification of both *EGFR* and *CDK4*. **d)** Heterogeneity of cells (measured by the Shannon index) across the E, C, EC and N/O (N/O = not amplified in either *CDK4* or *EGFR*) genotypes within each tumor. Each data point represents an image. Distinct colors represent different cores of the same tumor. **e)** Correlations between proportions of cells with distinct genotypes, with each point representing the ratio of the genotype within a tumor (i.e. consisting of all cells imaged within that tumor). Spearman correlation coefficients and p-values adjusted for multiple comparisons are shown along with best-fit curves. **f)** Within-core frequency of cells harboring TERT promoter mutation.

About half of all cells quantified in our samples were identified as N/O. This cell population could include cancer cells with distinct genotypes, without high copy number amplification of *EGFR* and *CDK4* genes but with other genetic alterations. It could also include cells of the tumor microenvironment that have a normal (non-amplified) genotype. Correlation of N/O cells frequency with DSP-based protein expression did not reveal a distinct identity of this population (**Supplemental Fig. 3b**). However, we found a negative correlation between the N/O cell frequency and expression of Olig2 (Spearman correlation −0.75, adjusted p-value 0.044). Our protein expression profiling found Olig2 expression to be strongly correlated with Sox2 expression (Spearman coefficient 0.81, adjusted p-value 0.007), although within the N/O population the negative correlation with Sox2 expression did not reach statistical significance. Both Olig2 and Sox2 are often expressed at high levels by GBM tumor cells (Verhaak et al., 2010; Q. Wang et al., 2017). Thus, low abundance of these proteins in N/O cells could indicate that a large proportion of these cells is of non-tumor origin.

Since cells with distinct genotypes based on the three selected amplifications were found in almost all tumors analyzed, we investigated whether there were any relationships between the proportions of these cell types in a tumor (**Fig. 2e, Supplemental Fig. 3c**). The prevalence of N/O cells in a tumor was negatively correlated with E cell frequency (Spearman correlation −0.91, adjusted p-value 1.7×10^−6^) and EC cell frequency (Spearman correlation −0.79, adjusted p-value 3.9×10^−4^), both of which were positively correlated with each other (Spearman correlation 0.78, adjusted p-value 5.7×10^−3^). EC cell frequency was also positively correlated, albeit more weakly, with C cell frequency (Spearman coefficient 0.66, adjusted p-value 5.7×10^−3^). No strong correlation was found between C and E cell frequency and a weak one was identified between C and N/O cell frequency (Spearman coefficient −0.52, adjusted p-value 0.04; **Supplemental Fig. 3c**).

Next, we sought to establish whether the tumor genotypes based on bulk tissue sequencing could be linked to diversity of single cell-based copy number-derived genotypes. Targeted sequencing of 50 genes using the OncoPanel (see **Methods**) showed only few mutations present in each tumor in our cohort and no clear distinction between tumors could be made based on these genetic alterations assessed in bulk (**Supplemental Fig. 3d, Supplemental Table 3**).

*hTERT* promoter hotspot mutations have been reported as one of the drivers of GBM (Körber et al., 2019; McKenna et al., 2013), yet they were not included in the targeted sequencing panel we used. To assess the frequency and spatial localization of cells harboring the hotspot mutation C228T in the *hTERT* promoter region, we performed *in situ* single cell mutation detection using the STAR-FISH assay (Janiszewska et al., 2015) on our multi-region GBM TMA (**Supplemental Fig. 4**). We found that the majority of the cores in the TMA contain over 50% TERT mutant cells, hetero- and/or homozygous (**Fig. 2f**). There was significant variation between distinct cores taken from different regions of the same tumor (**Supplemental Fig. 4d**). We did not observe any correlations between the ratio of cells with *hTERT* promoter mutation and the frequency of N/O cells, i.e. cells lacking *EGFR, CDK4*, and *PDGFRA* amplifications, most likely due to the heterogeneous nature of the N/O population (**Supplemental Fig. 4e**).

Together, these results show that single cell-based FISH quantification identifies a high prevalence of *hTERT* mutations and of cells harboring *CDK4, EGFR* and dual amplifications. These genetic alterations coexist in the majority of the GBM samples tested and exhibit a large extent of heterogeneity and distinct relationships in their frequencies.

### Single cell genotypes stratify tumors by co-occurrence of EGFR and CDK4 in the same cell versus distinct cell populations

Since amplifications of *EGFR, PDGFRA*, and *CDK4* were previously shown to correlate with the presence of transcriptionally distinct cell states in GBM (Neftel et al., 2019), cells harboring a combination of these genetic events could gain dual properties or have new distinct phenotypes. Our dataset of single cell genotypes allows for an investigation of differences between tumors in which these alterations tend to co-occur in the same cell and tumors in which they are more likely to be found in distinct cell populations. We therefore calculated the odds ratio of detecting co-amplified cells within a tumor for each of our 17 GBM cases. Since *PDGFRA* amplification was rare in our dataset, we again focused on *EGFR* and *CDK4* amplifications. The odds ratio (OR), as defined here, represents the likelihood that a cell harboring an *EGFR* amplification also harbors a *CDK4* amplification relative to the likelihood of a *CDK4* amplification unaccompanied by an *EGFR* amplification. At OR = 1, the presence of an amplification of one of these genes has no effect on the probability of the other gene being amplified. An OR above (below) unity therefore implies an increased (decreased) tendency for *EGFR* and *CDK4* amplifications to co-occur in the same cell over what would be expected if the presence of one amplification had no effect on the presence of the other. The OR calculation (see **Methods**) accounts for all relative frequencies of the cells with distinct genotypes (C, E, CE, N/O) within a tumor. Using the OR values, we separated the tumors into 3 classes based on OR tertiles (n=6, n=5, and n=6): OR^low^, where there is decreased tendency for *EGFR* and *CDK4* same-cell co-amplification (OR is significantly below unity); OR^high^, where there is increased tendency for same-cell co-amplification (OR is significantly above unity); and tumors where there is no strong effect either way (OR near unity) (**Fig. 3a,b, Supplemental Table 4**). Frequency of E cells (*EGFR* amplification alone) and N/O cells (lacking *EGFR* and *CDK4* amplifications) was significantly different between OR^high^ and OR^low^ groups, while C and EC cells were similarly distributed across all tumors (**Fig. 3c**). Thus, low OR can be driven by increased frequency of cells harboring only the *EGFR* amplification. While the median frequency of cells with *hTERT* promoter mutation was similar across the OR groups, OR^low^ tumors had significantly higher variability in *hTERT* mutation frequency (**Fig. 3c**). Interestingly, the average copy number of *EGFR* in E cells is significantly higher in OR^low^ tumors (**Fig. 3d**; mean 9.7 copies in OR^low^ vs 7.8 copies in OR^high^, Mann-Whitney test p-value 0.015) and also exhibits higher variability in OR^low^ tumors, as indicated by the Shannon diversity index (**Fig. 3f**). *CDK4* copy number diversity remained similar between OR^low^ and OR^high^ tumors (**Fig. 3e,f**).

**Figure 3.**
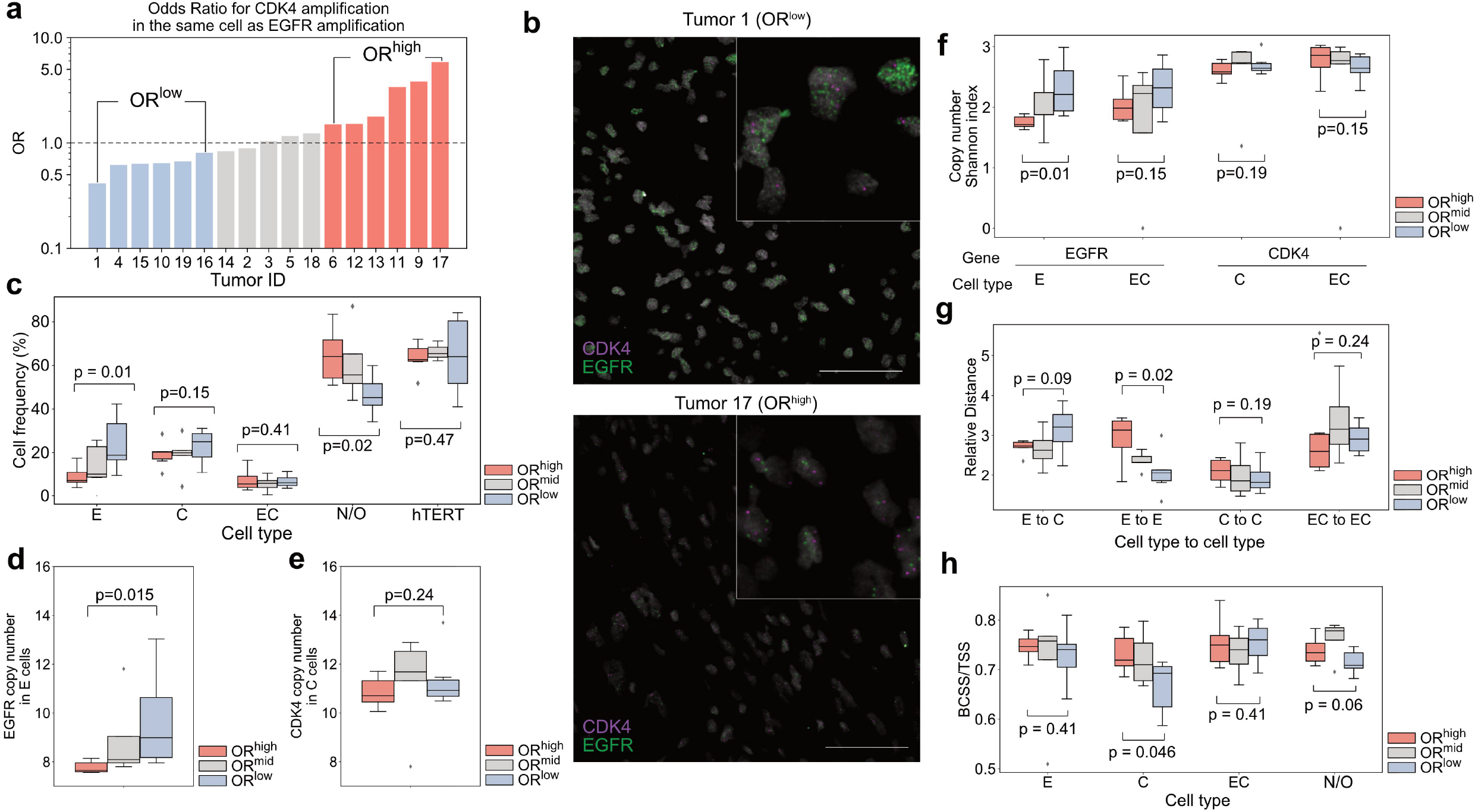
Single-cell genotypic classification of tumors based on relative proportions of cells with *EGFR* and *EGFR+CDK4* amplifications. **a)** Odds ratios for presence of *CDK4* amplification in the same cell as *EGFR* amplification, relative to cells with *CDK4* amplification alone. Odds ratio (OR) tertiles are shown in different colors. **b)** Representative images of *EGFR* and *CDK4* FISH in tumors with low (Tumor 1) and high (Tumor 17) odds ratio. Gray – DAPI, green – *EGFR* FISH, magenta – *CDK4* FISH. Scale bar – 50μm. **c)** Distribution of frequencies of E, C, EC, and N/O cells as well as cells harboring *TERT* promoter mutations in low- and high-OR tumors. P-values from a Mann-Whitney test of distributional differences are shown. **d)** Distributions of per-tumor mean *EGFR* copy number in E cells across the defined OR groups. **e)** Distributions of per-tumor mean *CDK4* copy number in C cells across the defined OR groups. **f)** Diversity (per-tumor Shannon index) of *EGFR* or *CDK4* copy number in single- and dual-amplified cells across the OR groups. **g)** Distributions of within-image distances between cells of different genotypes relative to distances between non-amplified-to-non-amplified cells in low- and high-OR tumors. **h)** Clustering of cells with distinct genotypes. BCSS/TSS represents the ratio of the between cluster sum of squares to the total sum of squares and provides a measure for the extent of clustering of each noted genotype, with a higher ratio indicating tighter clustering. All p-values shown in the above subfigures were obtained from Mann-Whitney tests. Colors of tumor groupings are indicated in (a). Genotypes are denoted as E – cells with amplified *EGFR* and non-amplified *CDK4*, C – cells with amplified CDK4 and non-amplified *EGFR*, EC – cells with amplification of both *EGFR* and *CDK4*, N/O – cells not amplified in either *EGFR* or *CDK4*. Each point within the displayed distributions represents a single tumor, with weighted averages (by cell number) employed for image-wise quantities (distances and clustering).

The *in situ* single-cell genotyping dataset contains spatial coordinates for each nucleus recorded in an image. This dataset thus enabled us to compute the distance between cells with distinct genotypes in OR^low^ and OR^high^ tumors. In comparison with OR^low^ tumors, E cells in OR^high^ tumors are farther away from each other (Mann-Whitney test p-value 0.02; **Fig. 3g**), while no significant difference in C cell distances was found between OR^low^ and OR^high^ tumors (distances shown are relative to N/O-N/O cell distances in order to normalize for variability in tissue density). Calculation of the between-cluster sum of squares to total sum of squares, which informs about the clustering of individual cell types, showed that in OR^high^ tumors C cells have a tendency toward tighter clustering within the tissue (Mann-Whitney test p-value 0.046, **Fig. 3h**).

Overall, our results suggest that the nature of *EGFR* amplification in E cells in tumors with lower prevalence of *CDK4* co-amplification may be qualitatively different from that observed in E cells in OR^high^ tumors. A possible explanation for the higher *EGFR* copy number frequency and diversity observed in OR^low^ tumors is that in these tumors *EGFR* amplification could be generated by extrachromosomal DNA fragments.

### Single cell EGFR-CDK4 amplification co-occurrence is associated with an immunosuppressive tumor microenvironment

Since the OR^low^ and OR^high^ tumor groups exhibit distinct cellular composition and distribution properties with respect to *EGFR* and *CDK4* copy number alterations, we sought to identify whether any proteins are differentially expressed between these tumor groups. We found that OR^high^ tumors were highly enriched in proteins associated with immune cell infiltration and an immunosuppressive microenvironment (CD163, IBA1, CD14, CD45, CD11b, HLA-DR; **Fig. 4a**), whereas these markers were downregulated in OR^low^ tumors. These markers additionally showed a trend towards a negative correlation with EGFR protein expression (Spearman correlation coefficient range −0.28 to −0.59; **Fig. 4b**), although these correlations were not statistically significant.

**Figure 4.**
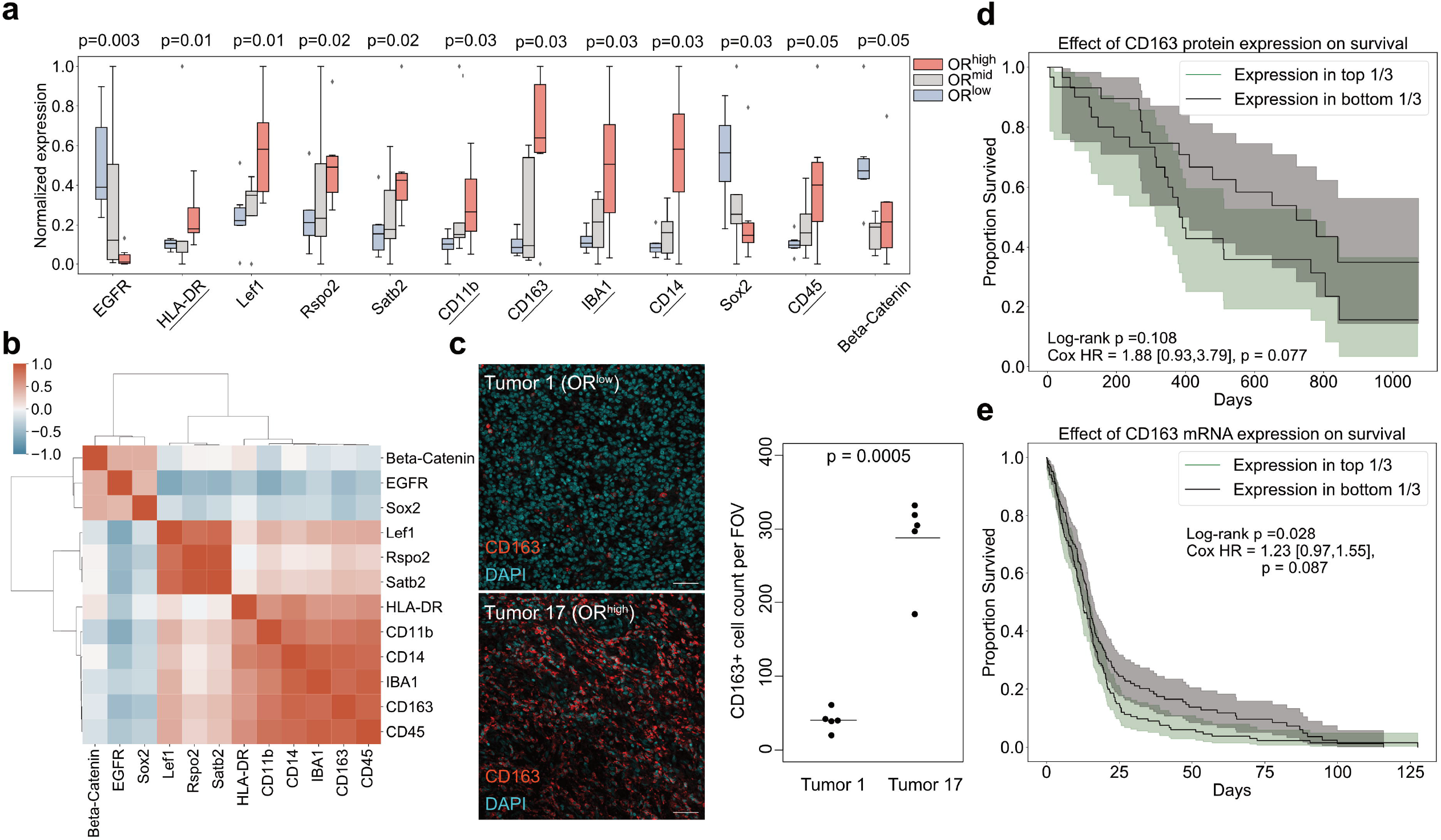
EGFR and CDK4 co-amplification odds ratio-based tumor groups have distinct protein expression patterns that are associated with immune infiltration and differential survival outcomes. **a)** Differential protein expression between odds ratio-based tumor groups. Each sample is a weighted mean of expression levels in individual ROIs in that tumor. Only proteins exhibiting a Mann-Whitney test p-value < 0.05 are shown (p-values shown above). Underlined proteins are markers of immune cells. **b)** Tumor-wise correlations between the differentially expressed proteins of panel a). **c)** Representative images of CD163 staining in the tumors with the highest and lowest co-amplification odds ratios in the dataset (left) and CD163+ cell quantification (right). FOV – field of view. Scale bar −50μm. **d)** Survival analysis performed on the CPTAC dataset (L.-B. Wang et al., 2021) for patients stratified into top and bottom tertiles (n=31 and n=30, respectively) based on CD163 protein level. Log-rank test p-values and Cox proportional hazard ratio (HR) values (age- and sex-adjusted, see **Methods**) are shown with a 95% confidence interval in brackets. Shaded areas represent confidence intervals. **e)** Survival analysis performed on the TCGA Firehose Legacy dataset for patients stratified into top and bottom tertiles (n=170 for each group) based on CD163 mRNA expression level. Log-rank test p-values and Cox proportional hazard ratio (HR) values (age- and sex-adjusted, see **Methods**) are shown with a 95% confidence interval in brackets. Shaded areas represent confidence intervals.

Among the other immune-related proteins in our expression panel, CD163, a marker of immunosuppressive macrophages, showed the highest mean difference in expression between the OR^high^ and OR^low^ tumor groups, suggesting enrichment of immunosuppressive cells (**Fig. 4a,b**). Immunofluorescent staining of the original sections from OR^low^ and OR^high^ tumors confirmed the striking difference in infiltration by CD163^+^ cells (**Fig. 4c**). Our results suggest that tumors classified based on the odds ratio of *EGFR* and *CDK4* co-occurrence at the single-cell level have different tumor microenvironments and that a higher relative frequency of co-amplification of these genes may promote a more immunosuppressive state. The presence of immunosuppressive macrophages and CD163 expression in the tumor has been previously linked with GBM patient survival (Liu et al., 2019; Zeng et al., 2020). In line with this, using CPTAC (L.-B. Wang et al., 2021) and TCGA (McKenna et al., 2013) datasets for CD163 protein and transcript level-based stratifications, respectively, we observed that GBM patients with higher CD163 expression tend to have somewhat shorter survival times, however in these datasets the differences are not significant (**Fig. 4d-e**).

## DISCUSSION

The insight provided by single cell studies into the diversity of genomic alterations can drive a deeper understanding of the subclonal structure of tumors not afforded by bulk tumor sequencing-based approaches. Key to this understanding is the interaction of a generally heterogeneous tumor microenvironment with subclonal-level changes. In this study, we aimed to gain a close-up view of this complex landscape through an integrative study of multi-region amplification and proteomic data in GBM biopsy samples. We identified distinct subclonal amplification patterns and associated microenvironmental properties and observed substantial spatial heterogeneity across all features that were profiled.

Central to these findings is our identification of the odds ratio of *EGFR*-*CDK4* co-amplification as an indicator of distinct alteration and expression trends. This quantity is a subclonal-level property that can be obtained from single cell data, but that is unavailable to bulk tumor sequencing studies. The odds ratio is based on the relative prevalence of distinct amplification-based cell types. High intratumor heterogeneity of these genetically-defined subpopulations may especially hamper the ability of bulk sequencing studies to identify distinct and prognostically-relevant patterns related to the extent of amplification occurrence and co-occurrence. This fact is exemplified in our study, as *EGFR* and *CDK4* amplifications (irrespective of PDGFRA amplification status) co-occur in 6.6% of nuclei analyzed, yet in 8 out of 17 of the tumors the frequency of these dual-amplified cells was more than three times higher than that in at least one imaged region. *PDGFRA* amplification was rare in our small dataset, and since our cohort is rich in *EGFR* amplified cases and a trend for mutual exclusivity between *EGFR* and *PDGFRA* amplification had previously been observed (Brennan et al., 2013; Szerlip et al., 2012), a larger cohort would likely be required to capture the relationships between *PDGFRA* amplification and the occurrence of *CDK4* and *EGFR* amplifications.

In our dataset, *EGFR*-amplified cells in tumors with a lower *EGFR*-*CDK4* co-amplification odds ratio had a higher average copy number and increased copy number diversity of *EGFR*. It is possible that *EGFR* amplification in extrachromosomal DNA (ecDNA) may explain these observations. ecDNA, circularized DNA fragments found in GBM and other tumors, contain an oncogene together with an enhancer element, allowing for increased gene expression (Dogan-Artun et al., 2019; Turner et al., 2017; Wu et al., 2019). The circularized form as well as lack of telomeres and centromeres contribute to highly efficient replication of ecDNA, which can then be asymmetrically divided between daughter cells in mitosis (Verhaak et al., 2019). This in turn can result in a large copy number of the oncogene and increased heterogeneity without the fitness penalty associated with large genomic alterations. In GBM, extrachromosomal amplification of *EGFR* is well documented (deCarvalho et al., 2018; Vogt et al., 2004). Thus, it is possible that our analysis identified tumors harboring ecDNA *EGFR* as a distinct class of tumors, associated with low immune marker expression and high levels of EGFR protein.

Our results demonstrate that higher relative frequencies of cells with co-amplification of *EGFR* and *CDK4* (or lower relative frequencies of cells harboring *EGFR* amplification alone) are associated with higher expression of immune markers, showing that OR^high^ tumors are likely highly infiltrated by immune cells. We found the strongest differential expression for CD163, a scavenger receptor expressed in macrophages and monocytes and a classical marker of M2/immunosuppressive polarization (Chen et al., 2019). This observation suggests that the immune cells infiltrating OR^high^ tumors may be predominantly of an immunosuppressive phenotype. The small size of our cohort (n=5-6 per group) precludes survival analysis; however, high expression of CD163 alone has been previously associated with poor outcome (Lisi et al., 2017; Liu et al., 2019; Zeng et al., 2020). Tumor infiltration by the immunosuppressive macrophages is a well-established hallmark of decreased survival for GBM patients (Quail and Joyce, 2017; Tomaszewski et al., 2019).

The link between GBM immune status and underlying genetic alterations in the tumor was explored in several studies. Most notably, inactivating mutations of the *NF1* gene were shown to be associated with increased expression of macrophage-related signatures (Luoto et al., 2018; Q. Wang et al., 2017). Tumor-infiltrating lymphocytes are depleted in tumors representing the classical-like subtype, harboring *EGFR* amplification and *PTEN* deletion (Rutledge et al., 2013). This finding is consistent with our observation that tumors with low *EGFR*-*CDK4* odds ratio, which are also enriched with cells with high *EGFR* copy number, had lower expression of immune markers. Interestingly, our data show that higher relative prevalence of cells with dual *CDK4* and *EGFR* amplification was associated with a macrophage-enriched tumor microenvironment. This effect cannot be explained by the presence of *CDK4*, as this amplification alone has been linked to lower macrophage infiltration and fewer CD4^+^ T cells in tumor bulk analysis (Luoto et al., 2018). Our findings therefore suggest that a subpopulation of GBM cells harboring a low level of *EGFR* amplification that co-occurs with amplification of *CDK4* is likely to elicit different immunological effects than subpopulations of cells with a single amplification. Further investigation of how these genetically distinct subpopulations interact with each other and with the cells of their native microenvironment will be necessary to exploit these properties as new therapeutic targets.

Our study suggests that the assessment of co-amplification of *EGFR* and *CDK4* at the single-cell level by FISH could serve as a proxy of immune status. Standard immunofluorescence to assess immune infiltration is often difficult to quantify due to variation in staining intensity driven by technical issues and imaging modality, and most notably due to limitations in accurate cell segmentation during image processing. Relying on a DNA-based nuclear signal of a FISH assay alleviates these challenges, as there is much less ambiguity about nuclear borders and speckle count compared to relative intensity of staining quantification. Quantification of FISH in thousands of cells could be easily implemented in a standard pathology lab setting, as our analyses were all done using open-source software and thresholding of the signal is easier to benchmark for FISH than for immunohistochemistry-based methods. Thus, the copy number alteration status of subclonal populations of cells within a tumor could serve as a representation of tumor-intrinsic properties that can be linked to tumor microenvironmental status. Future studies of larger cohorts, including samples from clinical trials targeting CKD4, EGFR or macrophages, will shed more light on the utility of single cell FISH signal quantification as a prognostic and predictive tool. Our study provides evidence that the presence of genetically distinct subpopulations is associated with differences in the tumor microenvironment.

## ACKNOWLEDGMENTS

We thank the members of Janiszewska and Michor laboratories for their critical reading of this manuscript and useful discussions. We thank Montina Van Meter from the Scripps Research Histology Core, Dr. Pabalu Karunadharma from Genomics Core, Dr. Adrian Reich from Bioinformatics Core for their dedication and technical expertise. We thank Dr. Liang Zhang from Nanostring for help with GeoMX DSP experiment. We thank Dr. Simona Cristea for useful discussions. This work was supported by the NIH K99/R00CA201606 (M.J.), the Center For Cancer Evolution (F.M.) and start-up funds from the Scripps Research Institute (M.J.).

## AUTHOR CONTRIBUTIONS

K.A.W. and D.E. are co-first authors, equally contributing to study conceptualization. K.A.W. and D.E. performed GeoMX data analysis. D.E. performed single-cell genotype analysis, integrative dataset analysis and survival data analysis. K.A.W. performed validation FISH and IHC staining. S.Y. performed TMA imaging and image analysis for FISH experiment. U.O. performed TMA imaging and image analysis for STAR-FISH experiment. R.S. optimized STAR-FISH TERT promoter mutation-specific conditions. C.V. provided clinical samples used to construct the TMA and evaluated histopathological features of the TMA H&E staining. T.O.M. consulted on the statistical analysis. F.M. and M.J. supervised the study. All authors helped design the study and write the manuscript.

## DECLARATION OF INTERESTS

M.J. is a member of a scientific advisory board at Viosera Therapeutics (ResistanceBio).

## SUPPLEMENTAL FIGURE LEGENDS

**Supplemental Figure 1. Spatial profiling ROI selection, clustering, protein expression and heterogeneity. a)** TMA morphology marker staining and ROI selection for digital spatial protein profiling. For cores with heterogeneous expression of CD45 (red), GFAP (green) and SMA (yellow), more than one ROI was selected. **b)** Unsupervised hierarchical clustering of ROIs based on expression level of 79 protein markers. Tumor IDs are color-coded.

**Supplemental Figure 2. Histopathology of TMA cores and correlations with protein expression**. **a)**Blood vessel quantification based on immunofluorescent ROI images. Vessel count was normalized to ROI area. Colors represent core of origin form each ROI. **b)**Representative image of Tumor 16 ROIs with low (left) and high (right) blood vessel count. Scale bar – 100μm. **c)**TMA core histology. Each bar represents a core, grouped by tumor ID. Tissue fraction occupied by tumor cells, normal brain, necrosis or hemorrhage was assessed based on H&E staining. **d)**Association between histopathological features in individual TMA cores and protein expression. Hierarchical clustering of core-level protein expression levels (average of ROI-level expression, weighted by number of nuclei in an ROI, for all ROIs in a core) shown alongside a histopathological categorization for each core. The scale bar represents the normalized expression level.

**Supplemental Figure 3. Validation of FISH and STAR-FISH assays. a)** FISH for *EGFR, CDK* and *PDGFRA* on histogel FFPE samples made from glioma cell lines with known amplification of the gene. Metaphase FISH on HEK293T cells is also shown. Scale bar −25 μm. **b)** Correlation of protein expression with non-amplified cell fraction for the 10 most strongly (negatively and positively) correlated proteins. **c)** Correlations between proportions of C (harboring *CDK4* amplification and no *EGFR* amplification) and N/O (no amplification of either *EGFR* or *CDK4*) cells and proportions of C and E (harboring *EGFR* amplification and no *CDK4* amplification) cells. Spearman correlation values and adjusted p-values are shown. **d)** Mutations identified by OncoPanel in 20 GBM tumors from which the TMA was derived. * - tumor 7 and 20 DNA extraction was successful, however the quality of the tissue on TMA was not sufficient for FISH analysis, thus these tumors were excluded from all TMA analyses. ClinVarID – clinical significance of the variants.

**Supplemental Figure 4. *In situ* single-cell *TERT* promoter mutation quantification. a)** Schematic of STAR-FISH assay for *TERT* promoter mutation detection *in situ*. 1^st^ round of the *in situ* PCR amplifies the *TERT* promoter region potentially harboring the mutation. In 2^nd^ round of the *in* situ PCR mismatched primers specifically amplify only the template harboring the C228T mutation, allowing for hybridization of a fluorescent probes to the overhangs within the PCR product. **b)** Validation of *TERT* promoter mutation PCR specificity on DNA extracted from wild-type (HEK293T) or mutant (U87) glioma cell lines. Arrows indicate the expected product size of 150bp. **c)** Validation of *TERT* promoter mutation-specific STAR-FISH, performed on intact U87 xenograft and a human GBM sample. U87 xenograft was subject to the assay in presence or absence of DNA polymerase (+/-Pol). Expected regional heterogeneity in human GBM sample was confirmed by digital droplet PCR results. Scale bar – 50μm.**d)** Proportion of cell with *TERT* promoter mutation in each tumor. Each datapoint represents an image.**e)** Correlations between proportions of cells with distinct genotypes. TERTp - *TERT* promoter mutation. E –cells with *EGFR* amplification only. C –cells with *CDK4* amplification only. EC –cells with *EGFR* and *CDK4* amplification. N/O – cells with no *EGFR* or *CDK4* amplification. Spearman correlation coefficients and p-values adjusted for multiple comparisons are shown.

## MATERIALS AND METHODS

### Human tissue samples

All experiments with use of human tumor tissue were approved by Scripps Research IRB protocol #IRB-18-7209 and Mount Sinai Medical Center IRB. Formalin-fixed paraffin embedded (FFPE) tissue blocks were provided by Dr. Cristina Vincentelli, Mount Sinai Medical Center, under IRB exemption for discarded tissue. GBM pathology was confirmed for each block by a board-certified neuropathologist. The cohort was comprised of 20 cases (10 female, 10 male) of GBM and one recurrence. Clinical details are shown in Supplemental Table 1. Tissue microarray (TMA) was constructed by manually selecting four distant areas within each block, cores of 1mm diameter were punched. H&E staining of the TMA was used to assess presence of hemorrhage, microvascular proliferations, and necrosis within each core, done by board certified pathologist, blinded to the other data derived from TMA.

### DNA sequencing

DNA was extracted from 30um sections of each FFPE GBM block, before cores for TMA construction were removed, using QiaAmp DNA FFPE Tissue Kit (Qiagen) and libraries were prepared using AmpliSeq Cancer HotSpot Panel v2 kit (Illumina). This panel targets 2800 COSMIC mutations in 50 oncogenes and tumor-suppressor genes. The libraries were sequenced using 2×150bp MiSeq format. For analysis DNA Amplicon Workflow version 3.24.1.8+master was used. Sequences were aligned to reference human genome hg19 using BWA-MEM Whole Genome Aligner version 0.7.9a-isis-1.0.2. Over 99.5% on-target aligned reads were reported for each sample, base mismatch was ~ 0.24%. Variant calling was done with Pisces Variant Caller 5.2.9.23, Illumina Annotation Engine 2.0.11-0-g7fb24a09, Bam Metrics v0.0.22, SAMtools 0.1.19-isis-1.0.3.

### GeoMx digital spatial profiling (DSP)

FFPE TMA was profiled using the commercial GeoMX DSP platform (Nanostring). TMA was stained with three fluorescent visualization markers, GFAP (GBM cells and astrocytes), CD45 (immune cells), and αSMA (endothelial cells) and two panels of UV-cleavable oligo-labeled antibodies (Suppl. Table XX). Stained slides were then digitally scanned, and 96 regions of interest (ROI) were selected based on the fluorescent channels, at least one ROI per TMA core. For TMA cores which displayed heterogeneity for the fluorescent markers, two ROIs were selected. Selected ROIs were UV-illuminated to release the conjugated oligos and quantified on the nCounter system (Nanostring).

### Fluorescence *in situ* hybridization (FISH)

Bacterial artificial chromosome clones RP11-339F13 (*EGFR* gene), RP11-231C18 (*PDGFRA* gene), RP11-571M6 (*CDK4* gene) were obtained from BACPAC Genomics (formerly BACPAC Resources at Childern’s Hospital Oakland Research Institute) and validated by PCR and FISH on xenografts with known amplification status (Supplemental Fig. 3). FISH probes were generated by Nick Translation (Abbott Molecular, 07J00-001) using fluorescent dUTPs (Abbott Molecular, 02N32-050). The FFPE TMA slide that was used in GeoMx DSP was de-mounted, washed in a series of increasing ethanol concentration solutions and incubated with hybridization solution containing 1:1:1 ratio of fluorescently labeled FISH probes, human COT1 DNA (Invitrogen, 15279011) and Vysis CEP hybridization buffer (Abbott Molecular, 07J36-001) at 74ºC for 7min and then at 37 ºC overnight. The slide was then washed for 2min with 0.4x SSC/0.3% NP-40 at room temperature, 0.4x SSC/0.3% NP-40 at 74ºC, 2x SSC/0.1% NP-40 at room temperature and 2x SSC at room temperature; buffers made with UltraPure 20x SSC (Invitrogen, 15557044) and NP-40 (Sigma, 13021). After wash with PBS and H_2_O, the slide was dried and mounted with ProLong Gold Antifade mountant with DAPI (Invitrogen, P36931). Imaging was performed on Olympus FV3000 confocal microscope (Olympus), for each TMA core three z-stack images were taken and maximum projections were used to quantify signals from the three fluorescent channels in each individual nucleus (ImageJ macro available upon request).

### STAR-FISH

The PCR conditions specific for *hTERT* C228T promoter mutation were optimized on DNA extracted from U87 (*hTERT* C228T mutant) and DBTRG cells (*hTERT* C228 WT) with primers F1: 5’CTATGGTTCCAGGC CCGTTC3’, R1: 5’GGCTCCCAGTGGATTCGC3’ in first round and F2: 5’TGTCGACGCAAAACCGGTTCCGGCCCAGCCCTTT3’, R2: 5’GCGATATGACGACGCGAATACCCACGTGCGCAGC3’ in second round (IDT). 1^st^ round PCR reactions were set with 2.5 mM MgCl_2_, 200 μM dNTPs (with 3:1 mix of 7-deaza-dGTP:dGTP; dNTPs from New England Bio Labs, cat#N0446, 7-deaza-2’-deoxy-GTP from Roche, cat#10988537001), 0.8 M Betaine (Alfa Aesar, cat#J77507VCR), 200 nM of each primer and 0.125 U of Platinum Taq Polymerase (Life Technologies, cat#10966083). After initial denaturation at 95 °C for 1 minute, 10 steps of 3 cycles each (95°C - 30 seconds, T_annealing_ - 30 seconds, 72°C - 20 seconds) were performed, starting at T_annealing_ = 70°C ending at T_annealing_ = 61°C with ΔT_annealing_ = −1°C between steps. Then 10 cycles were performed at T_annealing_ = 60°C (95°C - 30 seconds, 60°C - 30 seconds, 72°C - 20 seconds), followed by final extension at 72°C for 1 minute. PCR conditions for 2^nd^ round PCR were 2 mM MgCl_2_, 200 μM dNTPs (with 3:1 7-deaza-dGTP:dGTP), 250 nM of each primer, with 0.2 U of Platinum Taq Polymerase. After initial denaturation at 95 °C for 1 minute, 18 steps of 3 cycles each (95°C - 30 seconds, T_annealing_ - 30 seconds, 72°C - 20 seconds) were performed, starting at T_annealing_ = 70°C ending at T_annealing_ = 53°C with ΔT_annealing_ = −1°C between steps. Then 15 cycles were performed at T_annealing_ = 58°C (95°C - 30 seconds, 58°C - 30 seconds, 72°C - 20 seconds), followed by final extension at 72°C for 1 minute.

Specific-to-allele PCR-FISH was performed as described previously(Janiszewska et al., 2015). Briefly, after deparaffinization the FFPE TMA slide was treated with Proteinase K (20ug/ml, ThermoFisher AM2548) and subjected to two rounds of in situ PCR with a mixture of primers specific to the *hTERT* C228T mutation. After *in situ* PCR, the slide was washed in a series of increasing ethanol concentration solutions and hybridization of a probe specific to the PCR product (/5FAM/ +T+G+TCGACGCAAAACCGG+T+T+C (+ indicates LNA modified bases), custom made by Life Techonologies, working stock 25 μM) was performed at 74ºC for 7min and continued overnight at 37°C. After post-hybridization washes, as in FISH protocol, the slide was mounted with ProLong Gold Antifade mountant with DAPI (Invitrogen, P36931). Imaging, 3 images per core, and image analysis was performed as for the FISH experiment (modified ImageJ macro from our previous work (Janiszewska et al., 2021)). This quantification allowed for classification of each individual nucleus as WT or MUT for *hTERT* promoter mutation and downstream analysis was performed using R software.

### Immunofluorescent staining

After deparaffinization, the FFPE sections were subject to antigen retrieval solution pH9 (Dako, S2367) for 20 min in a steamer. Blocking with 10% goat serum (ThermoFisher, 31873) in PBST at room temperature was followed by incubation with primary antibody for CD163 (Abcam, ab182422, 1.4ug/ml working solution) at 4°C overnight. Next, the slides were washed 3 times with PBS and incubated for 1h at room temperature with goat anti-rabbit AlexaFluor-568 (ThermoFisher, Invitrogen A-1101, 5ug/ml working solution). After washes with PBS, the slides were mounted with with ProLong Gold Antifade mountant with DAPI (Invitrogen, P36931). Images were taken using Olympus FV3000 confocal microscope (Olympus) and image stitching was performed with CellSense (Olympus).

### Statistical analysis

#### GeoMX proteomic analysis

Tumor-level (core-level) protein expression was calculated from ROI-level protein expression as a weighted (by ROI nucleus number) average over all ROIs in a tumor (core) and min-max rescaled to a [0,1] range for each protein to enable global comparison. Tumor-level expression profiles were used in calculating Spearman correlations between proteins and performing hierarchical clustering on the dataset (Fig. 1c). ROI-level expression was used for establishing the relative inter-tumor to intra-tumor variability in protein expression via the Kruskal-Wallis H test (Python SciPy) (Fig. 1d). Here, each tumor constitutes a group, with within-group samples constituting all ROIs within a tumor. All p-values except where indicated are at the p < 0.01 significance level after FDR adjustment for multiple comparisons. Cluster-wise correlation between a cluster of immune markers and a cluster of neuronal markers (Spearman −0.48, p-value 0.052) was computed as the correlation between cluster scores comprising the sum of normalized protein levels for all proteins in the cluster, with distinct clusters indicated in Fig. 1c.

#### Single-cell genotype analysis

Cores for which no GeoMX data was available were removed from the analysis of both amplifications and *hTERT* mutations due to tissue quality concerns. Cells were considered to be amplified in a gene *g* if at least 6 copies of *g* were recorded in that nucleus location, analogous to the HER2 FISH scoring guidelines of the American Society of Clinical Oncology/College of American Pathologists (Li et al., 2020). Coordinates for which 50 or greater copies of any gene were recorded were removed from the dataset as these likely represent overlapping cells. The Shannon diversity index for genotypic diversity in each individual image *i* (Fig. 2d) was computed as *H*_*i*_ = −Σ_G∈ 𝒢_ *R*_*G,i*_ log *R*_*G,I*,_ where *R*_*G,I*,_ represents the proportion (if nonzero) of cells of genotype *G* in image *i* out of the set *𝒢* of four genotypes: E, C, EC, and N/O. Tumor-level genotype proportions were computed by combining data from all images in a particular tumor. Spearman correlations between proportions (Fig. 2e, Supp. Fig. 3c) were computed based on tumor-level proportions using the R Hmisc package, with p-values FDR-adjusted for multiple comparisons. Best-fit lines shown in Fig. 2e were obtained with an inverse (y = a/x + b) curve fit (Python SciPy) (top panels; negative Spearman coefficient) and linear fits (Python NumPy polyfit) (bottom panels; positive Spearman coefficient). Computation of minimum relative distances (Fig. 3g) was done by calculating in each image i, for each cell *c* of genotype *G* the Euclidean distance 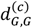 to the nearest cell of the same genotype and the Euclidean distance 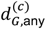 to the nearest cell of any genotype. The minimum relative distance for genotype *G* in image *i* is then given by

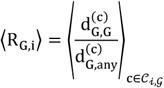

Where 𝒞_i,G_ is the set of all cells of genotype *G* in image *i* Computation of clustering properties proceeded by implementing k-means clustering (Python scikit-learn) in each image *i* for all cells *c* ∈ 𝒞_i,G_. The optimal number of clusters (from 2 to 5) was selected by finding the highest Silhouette score. The total sum of squares (TSS) was computed as Euclidean distances between all *c* ∈ 𝒞_i,G_. and the mean of *c, c* ∈ 𝒞_i, 𝒢_. The between cluster sum of squares (BCSS) was then computed as the difference between the TSS and the within cluster sum of squares, as given by the inertia property of the clusters obtained.

#### Integrative analysis

The odds ratio of *EGFR* and *CDK4* co-amplification (EC genotype) is given by the odds of a cell being co-amplified given that it is amplified in *EGFR* relative to the odds of a cell being co-amplified given that it is not amplified in *EGFR*, OR = (*N*_EC_/*N*_E_)/(*N*_C_/*N*_N/O_). The OR was computed in each tumor separately by combining data from all images in the tumor (Fig. 3a). For subsequent analysis, tumors were divided based on their computed OR into tertiles representing low-OR (OR^low^), middle-OR, and high-OR (OR^high^) tumors. To determine statistical significance of distributional differences between the top and bottom tertiles in copy numbers (Fig. 3a), distances (Fig. 3g), clustering (Fig. 3h), and protein expression (Fig. 4a) a Mann-Whitney test (Python SciPy) was employed. Associated proteomic analysis (Fig. 4a,b) employed tumor-level expression data, processed as described above.

#### Survival analysis

Separate analyses were carried out on clinical and proteomics data from Wang et al. (Cancer Cell 2021) and on clinical and mRNA expression levels (U133 microarray) TCGA Firehose Legacy dataset. In each case, clinical data was combined with CD163 expression data to fit a Cox proportional hazard model (Breslow’s method), adjusted for patient age and sex, and compute a 95% CI. IDH-mutant tumors were excluded from the analysis. The Python lifelines package was used to fit the model, obtain Kaplan-Meier curves, and perform a log-rank test.

## Notes

### Competing Interest Statement

M.J. is a member of a scientific advisory board at Viosera Therapeutics (ResistanceBio). All authors declare no competing interests.

## REFERENCES

Aldape, K., Brindle, K.M., Chesler, L., Chopra, R., Gajjar, A., Gilbert, M.R., Gottardo, N., Gutmann, D.H., Hargrave, D., Holland, E.C., Jones, D.T.W., Joyce, J.A., Kearns, P., Kieran, M.W., Mellinghoff, I.K., Merchant, M., Pfister, S.M., Pollard, S.M., Ramaswamy, V., Rich, J.N., Robinson, G.W., Rowitch, D.H., Sampson, J.H., Taylor, M.D., Workman, P., Gilbertson, R.J., 2019. Challenges to curing primary brain tumours. Nature Reviews Clinical Oncology 370, 699. doi:10.1038/s41571-019-0177-5

Barthel, F.P., Johnson, K.C., Varn, F.S., Moskalik, A.D., Tanner, G., Kocakavuk, E., Anderson, K.J., Abiola, O., Aldape, K., Alfaro, K.D., Alpar, D., Amin, S.B., Ashley, D.M., Bandopadhayay, P., Barnholtz-Sloan, J.S., Beroukhim, R., Bock, C., Brastianos, P.K., Brat, D.J., Brodbelt, A.R., Bruns, A.F., Bulsara, K.R., Chakrabarty, A., Chakravarti, A., Chuang, J.H., Claus, E.B., Cochran, E.J., Connelly, J., Costello, J.F., Finocchiaro, G., Fletcher, M.N., French, P.J., Gan, H.K., Gilbert, M.R., Gould, P.V., Grimmer, M.R., Iavarone, A., Ismail, A., Jenkinson, M.D., Khasraw, M., Kim, H., Kouwenhoven, M.C.M., LaViolette, P.S., Li, M., Lichter, P., Ligon, K.L., Lowman, A.K., Malta, T.M., Mazor, T., McDonald, K.L., Molinaro, A.M., Nam, D.-H., Nayyar, N., Ng, H.K., Ngan, C.Y., Niclou, S.P., Niers, J.M., Noushmehr, H., Noorbakhsh, J., Ormond, D.R., Park, C.-K., Poisson, L.M., Rabadan, R., Radlwimmer, B., Rao, G., Reifenberger, G., Sa, J.K., Schuster, M., Shaw, B.L., Short, S.C., Smitt, P.A.S., Sloan, A.E., Smits, M., Suzuki, H., Tabatabai, G., Van Meir, E.G., Watts, C., Weller, M., Wesseling, P., Westerman, B.A., Widhalm, G., Woehrer, A., Yung, W.K.A., Zadeh, G., Huse, J.T., De Groot, J.F., Stead, L.F., Verhaak, R.G.W., GLASS Consortium, 2019. Longitudinal molecular trajectories of diffuse glioma in adults. Nature 576, 112–120. doi:10.1038/s41586-019-1775-1

Brat, D.J., Castellano-Sanchez, A.A., Hunter, S.B., Pecot, M., Cohen, C., Hammond, E.H., Devi, S.N., Kaur, B., Van Meir, E.G., 2004. Pseudopalisades in glioblastoma are hypoxic, express extracellular matrix proteases, and are formed by an actively migrating cell population. Cancer Res 64, 920–927. doi:10.1158/0008-5472.can-03-2073

Brennan, C.W., Verhaak, R.G.W., McKenna, A., Campos, B., Noushmehr, H., Salama, S.R., Zheng, S., Chakravarty, D., Sanborn, J.Z., Berman, S.H., Beroukhim, R., Bernard, B., Wu, C.-J., Genovese, G., Shmulevich, I., Barnholtz-Sloan, J., Zou, L., Vegesna, R., Shukla, S.A., Ciriello, G., Yung, W.K., Zhang, W., Sougnez, C., Mikkelsen, T., Aldape, K., Bigner, D.D., Van Meir, E.G., Prados, M., Sloan, A., Black, K.L., Eschbacher, J., Finocchiaro, G., Friedman, W., Andrews, D.W., Guha, A., Iacocca, M., O’Neill, B.P., Foltz, G., Myers, J., Weisenberger, D.J., Penny, R., Kucherlapati, R., Perou, C.M., Hayes, D.N., Gibbs, R., Marra, M., Mills, G.B., Lander, E., Spellman, P., Wilson, R., Sander, C., Weinstein, J., Meyerson, M., Gabriel, S., Laird, P.W., Haussler, D., Getz, G., Chin, L., TCGA Research Network, 2013. The somatic genomic landscape of glioblastoma. Cell 155, 462–477. doi:10.1016/j.cell.2013.09.034

Calabrese, C., Poppleton, H., Kocak, M., Hogg, T.L., Fuller, C., Hamner, B., Oh, E.Y., Gaber, M.W., Finklestein, D., Allen, M., Frank, A., Bayazitov, I.T., Zakharenko, S.S., Gajjar, A., Davidoff, A., Gilbertson, R.J., 2007. A perivascular niche for brain tumor stem cells. Cancer Cell 11, 69–82. doi:10.1016/j.ccr.2006.11.020

Chen, T., Chen, J., Zhu, Y., Li, Y., Wang, Y., Chen, H., Wang, J., Li, X., Liu, Y., Li, B., Sun, X., Ke, Y., 2019. CD163, a novel therapeutic target, regulates the proliferation and stemness of glioma cells via casein kinase 2. Oncogene 38, 1183–1199. doi:10.1038/s41388-018-0515-6

Clevers, H., Loh, K.M., Nusse, R., 2014. Stem cell signaling. An integral program for tissue renewal and regeneration: Wnt signaling and stem cell control. Science 346, 1248012. doi:10.1126/science.1248012

deCarvalho, A.C., Kim, H., Poisson, L.M., Winn, M.E., Mueller, C., Cherba, D., Koeman, J., Seth, S., Protopopov, A., Felicella, M., Zheng, S., Multani, A., Jiang, Y., Zhang, J., Nam, D.-H., Petricoin, E.F., Chin, L., Mikkelsen, T., Verhaak, R.G.W., 2018. Discordant inheritance of chromosomal and extrachromosomal DNA elements contributes to dynamic disease evolution in glioblastoma. Nature Genetics 50, 708–717. doi:10.1038/s41588-018-0105-0

Dogan-Artun, N., Faber, Z.J., Bartels, C.F., Piazza, M.S., Allan, K.C., Mack, S.C., Gimple, R.C., Wu, Q., Rubin, B.P., Shetty, S., Sallari, R.C., Lupien, M., Rich, J.N., Scacheri, P.C., 2019. Functional Enhancers Shape Extrachromosomal Oncogene Amplifications. Cell 179, 1330–1341.e13. doi:10.1016/j.cell.2019.10.039

Garofano, L., Migliozzi, S., Oh, Y.T., D’Angelo, F., Najac, R.D., Ko, A., Frangaj, B., Caruso, F.P., Yu, K., Yuan, J., Zhao, W., Stefano, A.L.D., Bielle, F., Jiang, T., Sims, P., Suvà, M.L., Tang, F., Su, X.-D., Ceccarelli, M., Sanson, M., Lasorella, A., Iavarone, A., 2021. Pathway-based classification of glioblastoma uncovers a mitochondrial subtype with therapeutic vulnerabilities. Nature Cancer 2, 141–156. doi:10.1038/s43018-020-00159-4

Huang, M., Zhang, D., Wu, J.Y., Xing, K., Yeo, E., Li, C., Zhang, L., Holland, E., Yao, L., Qin, L., Binder, Z.A., O’Rourke, D.M., Brem, S., Koumenis, C., Gong, Y., Fan, Y., 2020. Wnt-mediated endothelial transformation into mesenchymal stem cell-like cells induces chemoresistance in glioblastoma. Sci Transl Med 12. doi:10.1126/scitranslmed.aay7522

Janiszewska, M., Liu, L., Almendro, V., Kuang, Y., Paweletz, C., Sakr, R.A., Weigelt, B., Hanker, A.B., Chandarlapaty, S., King, T.A., Reis-Filho, J.S., Arteaga, C.L., Park, S.Y., Michor, F., Polyak, K., 2015. In situ single-cell analysis identifies heterogeneity for PIK3CA mutation and HER2 amplification in HER2-positive breast cancer. Nature Genetics 47, 1212–1219. doi:10.1038/ng.3391

Janiszewska, M., Stein, S., Metzger Filho, O., Eng, J., Kingston, N.L., Harper, N.W., Rye, I.H., Alečković, M., Trinh, A., Murphy, K.C., Marangoni, E., Cristea, S., Oakes, B., Winer, E.P., Krop, I., Russnes, H.G., Spellman, P.T., Bucher, E., Hu, Z., Chin, K., Gray, J.W., Michor, F., Polyak, K., 2021. The impact of tumor epithelial and microenvironmental heterogeneity on treatment responses in HER2-positive breast cancer. JCI Insight 6. doi:10.1172/jci.insight.147617

Janiszewska, M., Suvà, M.L., Riggi, N., Houtkooper, R.H., Auwerx, J., Clément-Schatlo, V., Radovanovic, I., Rheinbay, E., Provero, P., Stamenkovic, I., 2012. Imp2 controls oxidative phosphorylation and is crucial for preserving glioblastoma cancer stem cells. Genes Dev. 26, 1926–1944. doi:10.1101/gad.188292.112

Körber, V., Yang, J., Barah, P., Wu, Y., Stichel, D., Gu, Z., Fletcher, M.N.C., Jones, D., Hentschel, B., Lamszus, K., Tonn, J.C., Schackert, G., Sabel, M., Felsberg, J., Zacher, A., Kaulich, K., Hübschmann, D., Herold-Mende, C., Deimling von, A., Weller, M., Radlwimmer, B., Schlesner, M., Reifenberger, G., Höfer, T., Lichter, P., 2019. Evolutionary Trajectories of IDHWT Glioblastomas Reveal a Common Path of Early Tumorigenesis Instigated Years ahead of Initial Diagnosis. Cancer Cell 35, 692–704.e12. doi:10.1016/j.ccell.2019.02.007

Lathia, J.D., Mack, S.C., Mulkearns-Hubert, E.E., Valentim, C.L.L., Rich, J.N., 2015. Cancer stem cells in glioblastoma. Genes Dev. 29, 1203–1217. doi:10.1101/gad.261982.115

Leder, K., Pitter, K., LaPlant, Q., Hambardzumyan, D., Ross, B.D., Chan, T.A., Holland, E.C., Michor, F., 2014. Mathematical modeling of PDGF-driven glioblastoma reveals optimized radiation dosing schedules. Cell 156, 603–616. doi:10.1016/j.cell.2013.12.029

Lee, J.-K., Wang, J., Sa, J.K., Ladewig, E., Lee, H.-O., Lee, I.-H., Kang, H.J., Rosenbloom, D.S., Camara, P.G., Liu, Z., van Nieuwenhuizen, P., Jung, S.W., Choi, S.W., Kim, J., Chen, A., Kim, K.-T., Shin, S., Seo, Y.J., Oh, J.-M., Shin, Y.J., Park, C.-K., Kong, D.-S., Seol, H.J., Blumberg, A., Lee, J.-I., Iavarone, A., Park, W.-Y., Rabadan, R., Nam, D.-H., 2017. Spatiotemporal genomic architecture informs precision oncology in glioblastoma. Nature Genetics 49, 594–599. doi:10.1038/ng.3806

Lee, J.H., Lee, J.E., Kahng, J.Y., Kim, S.H., Park, J.S., Yoon, S.J., Um, J.-Y., Kim, W.K., Lee, J.-K., Park, J., Kim, E.H., Lee, J.-H., Lee, J.-H., Chung, W.-S., Ju, Y.S., Park, S.-H., Chang, J.H., Kang, S.-G., Lee, J.H., 2018. Human glioblastoma arises from subventricular zone cells with low-level driver mutations. Nature 560, 243–247. doi:10.1038/s41586-018-0389-3

Li, A., Bai, Q., Kong, H., Zhou, S., Lv, H., Zhong, S., Li, M., Bi, R., Zhou, X., Yang, W., 2020. Impact of the Updated 2018 American Society of Clinical Oncology/College of American Pathologists Guideline for Human Epidermal Growth Factor Receptor 2 Testing in Breast Cancer. Arch Pathol Lab Med 144, 1097–1107. doi:10.5858/arpa.2019-0369-OA

Lisi, L., Ciotti, G.M.P., Braun, D., Kalinin, S., Currò, D., Russo Dello, C., Coli, A., Mangiola, A., Anile, C., Feinstein, D.L., Navarra, P., 2017. Expression of iNOS, CD163 and ARG-1 taken as M1 and M2 markers of microglial polarization in human glioblastoma and the surrounding normal parenchyma. Neurosci Lett 645, 106–112. doi:10.1016/j.neulet.2017.02.076

Liu, S., Zhang, C., Maimela, N.R., Yang, L., Zhang, Z., Ping, Y., Huang, L., Zhang, Y., 2019. Molecular and clinical characterization of CD163 expression via large-scale analysis in glioma. Oncoimmunology 8, 1601478. doi:10.1080/2162402X.2019.1601478

Luoto, S., Hermelo, I., Vuorinen, E.M., Hannus, P., Kesseli, J., Nykter, M., Granberg, K.J., 2018. Computational Characterization of Suppressive Immune Microenvironments in Glioblastoma. Cancer Res 78, 5574–5585. doi:10.1158/0008-5472.CAN-17-3714

Magurran, A.E., 2005. Biological diversity. Curr Biol 15, R116–8. doi:10.1016/j.cub.2005.02.006

McKenna, A., Campos, B., Salama, S.R., Zheng, S., Chakravarty, D., Sanborn, J.Z., Berman, S.H., Bernard, B., Wu, C.-J., Genovese, G., Shmulevich, I., Barnholtz-Sloan, J., Zou, L., Vegesna, R., Shukla, S.A., Ciriello, G., Yung, W.K., Zhang, W., Bigner, D.D., Van Meir, E.G., Prados, M., Sloan, A., Black, K.L., Eschbacher, J., Friedman, W., Andrews, D.W., Guha, A., Iacocca, M., O’Neill, B.P., Foltz, G., Myers, J., Weisenberger, D.J., Penny, R., Kucherlapati, R., Gibbs, R., Marra, M., Mills, G.B., Lander, E., Spellman, P., Wilson, R., Sander, C., Weinstein, J., Meyerson, M., Gabriel, S., Laird, P.W., Haussler, D., Getz, G., Benz, C., Barnholtz-Sloan, J., Barrett, W., Ostrom, Q., Wolinsky, Y., Black, K.L., Bose, B., Boulos, P.T., Boulos, M., Brown, J., Czerinski, C., Eppley, M., Iacocca, M., Kempista, T., Kitko, T., Koyfman, Y., Rabeno, B., Rastogi, P., Sugarman, M., Swanson, P., Yalamanchii, K., Otey, I.P., Liu, Y.S., Xiao, Y., Auman, J.T., Chen, P.-C., Hadjipanayis, A., Lee, E., Lee, S., Park, P.J., Seidman, J., Yang, L., Kucherlapati, R., Kalkanis, S., Mikkelsen, T., Poisson, L.M., Raghunathan, A., Scarpace, L., Bernard, B., Bressler, R., Eakin, A., Iype, L., Kreisberg, R.B., Leinonen, K., Reynolds, S., Rovira, H., Thorsson, V., Shmulevich, I., Annala, M.J., Penny, R., Paulauskis, J., Curley, E., Hatfield, M., Mallery, D., Morris, S., Shelton, T., Shelton, C., Sherman, M., Yena, P., Cuppini, L., DiMeco, F., Eoli, M., Finocchiaro, G., Maderna, E., Pollo, B., Saini, M., Balu, S., Hoadley, K.A., Li, L., Miller, C.R., Shi, Y., Topal, M.D., Wu, J., Dunn, G., Giannini, C., O’Neill, B.P., Aksoy, B.A., Antipin, Y., Borsu, L., Berman, S.H., Brennan, C.W., Cerami, E., Chakravarty, D., Gao, J., Gross, B., Jacobsen, A., Ladanyi, M., Lash, A., Liang, Y., Reva, B., Schultz, N., Shen, R., Socci, N.D., Viale, A., Ferguson, M.L., Chen, Q.-R., Demchok, J.A., Dillon, L.A.L., Shaw, K.R.M., Sheth, M., Tarnuzzer, R., Wang, Z., Yang, L., Davidsen, T., Guyer, M.S., Ozenberger, B.A., Sofia, H.J., Bergsten, J., Eckman, J., Harr, J., Myers, J., Smith, C., Tucker, K., Winemiller, C., Zach, L.A., Ljubimova, J.Y., Eley, G., Ayala, B., Jensen, M.A., Kahn, A., Pihl, T.D., Pot, D.A., Wan, Y., Eschbacher, J., Foltz, G., Hansen, N., Hothi, P., Lin, B., Shah, N., Yoon, J.-G., Lau, C., Berens, M., Ardlie, K., Beroukhim, R., Carter, S.L., Cherniack, A.D., Noble, M., Cho, J., Cibulskis, K., DiCara, D., Frazer, S., Gabriel, S.B., Gehlenborg, N., Gentry, J., Heiman, D., Kim, J., Jing, R., Lander, E.S., Lawrence, M., Lin, P., Mallard, W., Onofrio, R.C., Saksena, G., Schumacher, S., Sougnez, C., Stojanov, P., Tabak, B., Voet, D., Zhang, H., Dees, N.N., Ding, L., Fulton, L.L., Fulton, R.S., Kanchi, K.-L., Mardis, E.R., Wilson, R.K., Baylin, S.B., Andrews, D.W., Harshyne, L., Cohen, M.L., Devine, K., Sloan, A.E., VandenBerg, S.R., Berger, M.S., Prados, M., Carlin, D., Craft, B., Ellrott, K., Goldman, M., Goldstein, T., Grifford, M., Haussler, D., Ma, S., Ng, S., Salama, S.R., Sanborn, J.Z., Stuart, J., Swatloski, T., Waltman, P., Zhu, J., Foss, R., Frentzen, B., Friedman, W., McTiernan, R., Yachnis, A., Hayes, D.N., Perou, C.M., Vegesna, R., Mao, Y., Akbani, R., Aldape, K., Bogler, O., Fuller, G.N., Liu, W., Liu, Y., Lu, Y., Mills, G., Protopopov, A., Ren, X., Sun, Y., Wu, C.-J., Yung, W.K.A., Zhang, W., Zhang, J., Chen, K., Weinstein, J.N., Chin, L., Verhaak, R.G.W., Noushmehr, H., Weisenberger, D.J., Bootwalla, M.S., Lai, P.H., Triche, T.J., Jr., Van Den Berg, D.J., Gutmann, D.H., Lehman, N.L., VanMeir, E.G., Brat, D., Olson, J.J., Mastrogianakis, G.M., Devi, N.S., Zhang, Z., Bigner, D., Lipp, E., McLendon, R., 2013. The Somatic Genomic Landscape of Glioblastoma. Cell 155, 462–477. doi:10.1016/j.cell.2013.09.034

Neftel, C., Laffy, J., Filbin, M.G., Hara, T., Shore, M.E., Rahme, G.J., Richman, A.R., Silverbush, D., Shaw, M.L., Hebert, C.M., Dewitt, J., Gritsch, S., Perez, E.M., Gonzalez Castro, L.N., Lan, X., Druck, N., Rodman, C., Dionne, D., Kaplan, A., Bertalan, M.S., Small, J., Pelton, K., Becker, S., Bonal, D., Nguyen, Q.-D., Servis, R.L., Fung, J.M., Mylvaganam, R., Mayr, L., Gojo, J., Haberler, C., Geyeregger, R., Czech, T., Slavc, I., Nahed, B.V., Curry, W.T., Carter, B.S., Wakimoto, H., Brastianos, P.K., Batchelor, T.T., Stemmer-Rachamimov, A., Martinez-Lage, M., Frosch, M.P., Stamenkovic, I., Riggi, N., Rheinbay, E., Monje, M., Rozenblatt-Rosen, O., Cahill, D.P., Patel, A.P., Hunter, T., Verma, I.M., Ligon, K.L., Louis, D.N., Regev, A., Bernstein, B.E., Tirosh, I., Suvà, M.L., 2019. An Integrative Model of Cellular States, Plasticity, and Genetics for Glioblastoma. Cell 178, 835–849.e21. doi:10.1016/j.cell.2019.06.024

Nicholson, J.G., Fine, H.A., 2021. Diffuse Glioma Heterogeneity and Its Therapeutic Implications. Cancer Discov 11, 575–590. doi:10.1158/2159-8290.CD-20-1474

O’Connor, J.P.B., Rose, C.J., Waterton, J.C., Carano, R.A.D., Parker, G.J.M., Jackson, A., 2015. Imaging intratumor heterogeneity: role in therapy response, resistance, and clinical outcome. Clin. Cancer Res. 21, 249–257. doi:10.1158/1078-0432.CCR-14-0990

Ozawa, T., Riester, M., Cheng, Y.-K., Huse, J.T., Squatrito, M., Helmy, K., Charles, N., Michor, F., Holland, E.C., 2014. Most human non-GCIMP glioblastoma subtypes evolve from a common proneural-like precursor glioma. Cancer Cell 26, 288–300. doi:10.1016/j.ccr.2014.06.005

Patel, A.P., Tirosh, I., Trombetta, J.J., Shalek, A.K., Gillespie, S.M., Wakimoto, H., Cahill, D.P., Nahed, B.V., Curry, W.T., Martuza, R.L., Louis, D.N., Rozenblatt-Rosen, O., Suvà, M.L., Regev, A., Bernstein, B.E., 2014. Single-cell RNA-seq highlights intratumoral heterogeneity in primary glioblastoma. Science 344, 1396–1401. doi:10.1126/science.1254257

Prager, B.C., Xie, Q., Bao, S., Rich, J.N., 2019. Cancer Stem Cells: The Architects of the Tumor Ecosystem. Cell Stem Cell 24, 41–53. doi:10.1016/j.stem.2018.12.009

Rheinbay, E., Suvà, M.L., Gillespie, S.M., Wakimoto, H., Patel, A.P., Shahid, M., Oksuz, O., Rabkin, S.D., Martuza, R.L., Rivera, M.N., Louis, D.N., Kasif, S., Chi, A.S., Bernstein, B.E., 2013. An aberrant transcription factor network essential for Wnt signaling and stem cell maintenance in glioblastoma. Cell Reports 3, 1567–1579. doi:10.1016/j.celrep.2013.04.021

Rich, J.N., Reardon, D.A., Peery, T., Dowell, J.M., Quinn, J.A., Penne, K.L., Wikstrand, C.J., Van Duyn, L.B., Dancey, J.E., McLendon, R.E., Kao, J.C., Stenzel, T.T., Ahmed Rasheed, B.K., Tourt-Uhlig, S.E., Herndon, J.E., Vredenburgh, J.J., Sampson, J.H., Friedman, A.H., Bigner, D.D., Friedman, H.S., 2004. Phase II trial of gefitinib in recurrent glioblastoma. J Clin Oncol 22, 133–142. doi:10.1200/JCO.2004.08.110

Rong, Y., Durden, D.L., Van Meir, E.G., Brat, D.J., 2006. “Pseudopalisading” necrosis in glioblastoma: a familiar morphologic feature that links vascular pathology, hypoxia, and angiogenesis. J. Neuropathol. Exp. Neurol. 65, 529–539. doi:10.1097/00005072-200606000-00001

Rutledge, W.C., Kong, J., Gao, J., Gutman, D.A., Cooper, L.A.D., Appin, C., Park, Y., Scarpace, L., Mikkelsen, T., Cohen, M.L., Aldape, K.D., McLendon, R.E., Lehman, N.L., Miller, C.R., Schniederjan, M.J., Brennan, C.W., Saltz, J.H., Moreno, C.S., Brat, D.J., 2013. Tumor-infiltrating lymphocytes in glioblastoma are associated with specific genomic alterations and related to transcriptional class. Clin. Cancer Res. 19, 4951–4960. doi:10.1158/1078-0432.CCR-13-0551

Snuderl, M., Fazlollahi, L., Le, L.P., Nitta, M., Zhelyazkova, B.H., Davidson, C.J., Akhavanfard, S., Cahill, D.P., Aldape, K.D., Betensky, R.A., Louis, D.N., Iafrate, A.J., 2011. Mosaic amplification of multiple receptor tyrosine kinase genes in glioblastoma. Cancer Cell 20, 810–817. doi:10.1016/j.ccr.2011.11.005

Sottoriva, A., Spiteri, I., Piccirillo, S.G.M., Touloumis, A., Collins, V.P., Marioni, J.C., Curtis, C., Watts, C., Tavaré, S., 2013. Intratumor heterogeneity in human glioblastoma reflects cancer evolutionary dynamics. Proc. Natl. Acad. Sci. U.S.A. 110, 4009–4014. doi:10.1073/pnas.1219747110

Szerlip, N.J., Pedraza, A., Chakravarty, D., Azim, M., McGuire, J., Fang, Y., Ozawa, T., Holland, E.C., Huse, J.T., Jhanwar, S., Leversha, M.A., Mikkelsen, T., Brennan, C.W., 2012. Intratumoral heterogeneity of receptor tyrosine kinases EGFR and PDGFRA amplification in glioblastoma defines subpopulations with distinct growth factor response. Proc. Natl. Acad. Sci. U.S.A. 109, 3041–3046. doi:10.1073/pnas.1114033109

Tehrani, M., Friedman, T.M., Olson, J.J., Brat, D.J., 2008. Intravascular thrombosis in central nervous system malignancies: a potential role in astrocytoma progression to glioblastoma. Brain Pathol 18, 164–171. doi:10.1111/j.1750-3639.2007.00108.x

Tomaszewski, W., Sanchez-Perez, L., Gajewski, T.F., Sampson, J.H., 2019. Brain Tumor Microenvironment and Host State: Implications for Immunotherapy. Clin. Cancer Res. 25, 4202–4210. doi:10.1158/1078-0432.CCR-18-1627

Turner, K.M., Deshpande, V., Beyter, D., Koga, T., Rusert, J., Lee, C., Bin Li Arden, K., Ren, B., Nathanson, D.A., Kornblum, H.I., Taylor, M.D., Kaushal, S., Cavenee, W.K., Wechsler-Reya, R., Furnari, F.B., VandenBerg, S.R., Rao, P.N., Wahl, G.M., Bafna, V., Mischel, P.S., 2017. Extrachromosomal oncogene amplification drives tumour evolution and genetic heterogeneity. Nature 543, 122–125. doi:10.1038/nature21356

Verhaak, R.G.W., Bafna, V., Mischel, P.S., 2019. Extrachromosomal oncogene amplification in tumour pathogenesis and evolution. Nature Reviews Cancer 19, 283–288. doi:10.1038/s41568-019-0128-6

Verhaak, R.G.W., Hoadley, K.A., Purdom, E., Wang, V., Qi, Y., Wilkerson, M.D., Miller, C.R., Ding, L., Golub, T., Mesirov, J.P., Alexe, G., Lawrence, M., O’Kelly, M., Tamayo, P., Weir, B.A., Gabriel, S., Winckler, W., Gupta, S., Jakkula, L., Feiler, H.S., Hodgson, J.G., James, C.D., Sarkaria, J.N., Brennan, C., Kahn, A., Spellman, P.T., Wilson, R.K., Speed, T.P., Gray, J.W., Meyerson, M., Getz, G., Perou, C.M., Hayes, D.N., Cancer Genome Atlas Research Network, 2010. Integrated genomic analysis identifies clinically relevant subtypes of glioblastoma characterized by abnormalities in PDGFRA, IDH1, EGFR, and NF1. Cancer Cell 17, 98–110. doi:10.1016/j.ccr.2009.12.020

Vogt, N., Lefèvre, S.-H., Apiou, F., Dutrillaux, A.-M., Cör, A., Leuraud, P., Poupon, M.-F., Dutrillaux, B., Debatisse, M., Malfoy, B., 2004. Molecular structure of double-minute chromosomes bearing amplified copies of the epidermal growth factor receptor gene in gliomas. PNAS 101, 11368–11373. doi:10.1073/pnas.0402979101

Wang, L.-B., Karpova, A., Gritsenko, M.A., Kyle, J.E., Cao, S., Li, Y., Rykunov, D., Colaprico, A., Rothstein, J.H., Hong, R., Stathias, V., Cornwell, M., Petralia, F., Wu, Y., Reva, B., Krug, K., Pietro Pugliese, Kawaler, E., Olsen, L.K., Liang, W.-W., Song, X., Dou, Y., Wendl, M.C., Caravan, W., Liu, W., Zhou, D.C., Ji, J., Tsai, C.-F., Petyuk, V.A., Moon, J., Ma, W., Chu, R.K., Weitz, K.K., Moore, R.J., Monroe, M.E., Zhao, R., Yang, X., Yoo, S., Krek, A., Demopoulos, A., Zhu, H., Wyczalkowski, M.A., McMichael, J.F., Henderson, B.L., Lindgren, C.M., Boekweg, H., Lu, S., Baral, J., Yao, L., Stratton, K.G., Bramer, L.M., Zink, E., Couvillion, S.P., Bloodsworth, K.J., Satpathy, S., Sieh, W., Boca, S.M., Schürer, S., Chen, F., Wiznerowicz, M., Ketchum, K.A., Boja, E.S., Kinsinger, C.R., Robles, A.I., Hiltke, T., Thiagarajan, M., Nesvizhskii, A.I., Zhang, B., Mani, D.R., Ceccarelli, M., Chen, X.S., Cottingham, S.L., Li, Q.K., Kim, A.H., Fenyö, D., Ruggles, K.V., Rodriguez, H., Mesri, M., Payne, S.H., Resnick, A.C., Wang, P., Smith, R.D., Iavarone, A., Chheda, M.G., Barnholtz-Sloan, J.S., Rodland, K.D., Liu, T., Ding, L., Consortium, C.P.T.A., Agarwal, A., Amin, M., An, E., Anderson, M.L., Andrews, D.W., Bauer, T., Birger, C., Birrer, M.J., Blumenberg, L., Bocik, W.E., Borate, U., Borucki, M., Burke, M.C., Cai, S., Calinawan, A.P., Carr, S.A., Cerda, S., Chan, D.W., Charamut, A., Chen, L.S., Chesla, D., Chinnaiyan, A.M., Chowdhury, S., Cieślik, M.P., Clark, D.J., Culpepper, H., Czernicki, T., D’Angelo, F., Day, J., De Young, S., Demir, E., Dhanasekaran, S.M., Dhir, R., Domagalski, M.J., Druker, B., Duffy, E., Dyer, M., Edwards, N.J., Edwards, R., Elburn, K., Ellis, M.J., Eschbacher, J., Francis, A., Gabriel, S., Gabrovski, N., Garofano, L., Getz, G., Gillette, M.A., Godwin, A.K., Golbin, D., Hanhan, Z., Hannick, L.I., Hariharan, P., Hindenach, B., Hoadley, K.A., Hostetter, G., Huang, C., Jaehnig, E., Jewell, S.D., Ji, N., Jones, C.D., Karz, A., Kaspera, W., Kim, L., Kothadia, R.B., Kumar-Sinha, C., Lei, J., Leprevost, F.D., Li, K., Liao, Y., Lilly, J., Liu, H., Lubínski, J., Madan, R., Maggio, W., Malc, E., Malovannaya, A., Mareedu, S., Markey, S.P., Marrero-Oliveras, A., Martinez, N., Maunganidze, N., McDermott, J.E., McGarvey, P.B., McGee, J., Mieczkowski, P., Migliozzi, S., Modugno, F., Montgomery, R., Newton, C.J., Omenn, G.S., Ozbek, U., Paklina, O.V., Paulovich, A.G., Perou, A.M., Pico, A.R., Piehowski, P.D., Placantonakis, D.G., Polonskaya, L., Potapova, O., Pruetz, B., Qi, L., Ramkissoon, S., Resnick, A., Richey, S., Riggins, G., Robinson, K., Roche, N., Rohrer, D.C., Rood, B.R., Rossell, L., Savage, S.R., Schadt, E.E., Shi, Y., Shi, Z., Shutack, Y., Singh, S., Skelly, T., Sokoll, L.J., Stawicki, J., Stein, S.E., Suh, J., Szopa, W., Tabor, D., Tan, D., Tansil, D., Thangudu, R.R., Tognon, C., Traer, E., Tsang, S., Tyner, J., Um, K.S., Valley, D.R., Vasaikar, S., Vatanian, N., Velvulou, U., Vernon, M., Wan, W., Wang, J., Webster, A., Wen, B., Whiteaker, J.R., Wilson, G.D., Zakhartsev, Y., Zelt, R., Zhang, H., Zhang, L., Zhang, Z., Zhao, G., Zhu, J., 2021. Proteogenomic and metabolomic characterization of human glioblastoma. Cancer Cell 39, 509–527.e21. doi:10.1016/j.ccell.2021.01.006

Wang, Q., Hu, B., Hu, X., Kim, H., Squatrito, M., Scarpace, L., deCarvalho, A.C., Lyu, S., Li, P., Li, Y., Barthel, F., Cho, H.J., Lin, Y.-H., Satani, N., Martinez-Ledesma, E., Zheng, S., Chang, E., Sauvé, C.-E.G., Olar, A., Lan, Z.D., Finocchiaro, G., Phillips, J.J., Berger, M.S., Gabrusiewicz, K.R., Wang, G., Eskilsson, E., Hu, J., Mikkelsen, T., DePinho, R.A., Muller, F., Heimberger, A.B., Sulman, E.P., Nam, D.-H., Verhaak, R.G.W., 2017. Tumor Evolution of Glioma-Intrinsic Gene Expression Subtypes Associates with Immunological Changes in the Microenvironment. Cancer Cell 32, 42–56.e6. doi:10.1016/j.ccell.2017.06.003

Wang, Z., Sun, D., Chen, Y.-J., Xie, X., Shi, Y., Tabar, V., Brennan, C.W., Bale, T.A., Jayewickreme, C.D., Laks, D.R., Alcantara Llaguno, S., Parada, L.F., 2020. Cell Lineage-Based Stratification for Glioblastoma. Cancer Cell. doi:10.1016/j.ccell.2020.06.003

Wen, P.Y., Chang, S.M., Van den Bent, M.J., Vogelbaum, M.A., Macdonald, D.R., Lee, E.Q., 2017. Response Assessment in Neuro-Oncology Clinical Trials. J Clin Oncol 35, 2439–2449. doi:10.1200/JCO.2017.72.7511

Wen, P.Y., Weller, M., Lee, E.Q., Alexander, B.M., Barnholtz-Sloan, J.S., Barthel, F.P., Batchelor, T.T., Bindra, R.S., Chang, S.M., Chiocca, E.A., Cloughesy, T.F., DeGroot, J.F., Galanis, E., Gilbert, M.R., Hegi, M.E., Horbinski, C., Huang, R.Y., Lassman, A.B., Le Rhun, E., Lim, M., Mehta, M.P., Mellinghoff, I.K., Minniti, G., Nathanson, D., Platten, M., Preusser, M., Roth, P., Sanson, M., Schiff, D., Short, S.C., Taphoorn, M.J.B., Tonn, J.-C., Tsang, J., Verhaak, R.G.W., Deimling von, A., Wick, W., Zadeh, G., Reardon, D.A., Aldape, K.D., Van den Bent, M.J., 2020. Glioblastoma in adults: a Society for Neuro-Oncology (SNO) and European Society of Neuro-Oncology (EANO) consensus review on current management and future directions. Neuro-Oncology 22, 1073–1113. doi:10.1093/neuonc/noaa106

Wen, P.Y., Yung, W.K.A., Lamborn, K.R., Dahia, P.L., Wang, Y., Peng, B., Abrey, L.E., Raizer, J., Cloughesy, T.F., Fink, K., Gilbert, M., Chang, S., Junck, L., Schiff, D., Lieberman, F., Fine, H.A., Mehta, M., Robins, H.I., DeAngelis, L.M., Groves, M.D., Puduvalli, V.K., Levin, V., Conrad, C., Maher, E.A., Aldape, K., Hayes, M., Letvak, L., Egorin, M.J., Capdeville, R., Kaplan, R., Murgo, A.J., Stiles, C., Prados, M.D., 2006. Phase I/II study of imatinib mesylate for recurrent malignant gliomas: North American Brain Tumor Consortium Study 99-08. Clin. Cancer Res. 12, 4899–4907. doi:10.1158/1078-0432.CCR-06-0773

Wu, S., Turner, K.M., Nguyen, N., Raviram, R., Erb, M., Santini, J., Luebeck, J., Rajkumar, U., Diao, Y., Li, B., Zhang, W., Jameson, N., Corces, M.R., Granja, J.M., Chen, X., Coruh, C., Abnousi, A., Houston, J., Ye, Z., Hu, R., Yu, M., Kim, H., Law, J.A., Verhaak, R.G.W., Hu, M., Furnari, F.B., Chang, H.Y., Ren, B., Bafna, V., Mischel, P.S., 2019. Circular ecDNA promotes accessible chromatin and high oncogene expression. Nature 575, 699–703. doi:10.1038/s41586-019-1763-5

Zeng, F., Wang, K., Liu, X., Zhao, Z., 2020. Comprehensive profiling identifies a novel signature with robust predictive value and reveals the potential drug resistance mechanism in glioma. Cell Commun Signal 18, 2–13. doi:10.1186/s12964-019-0492-6

