## Supplementary figures and images for "Single-cell genetic heterogeneity linked to immune infiltration in glioblastoma"

### Supplemental Figures

#### Supplemental Fig. 1

a

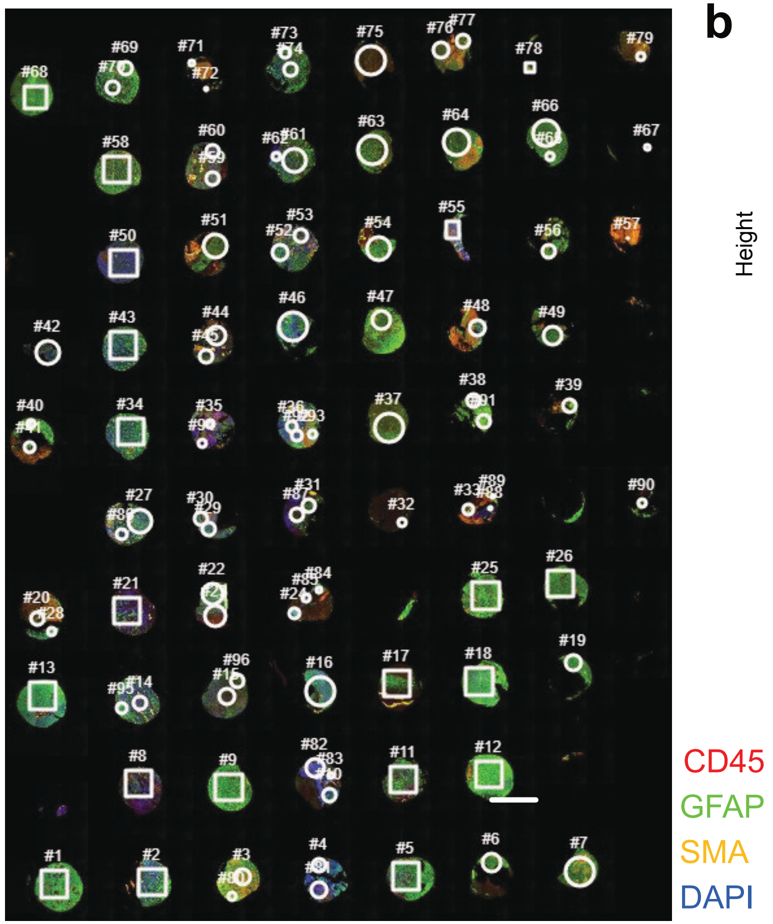


b

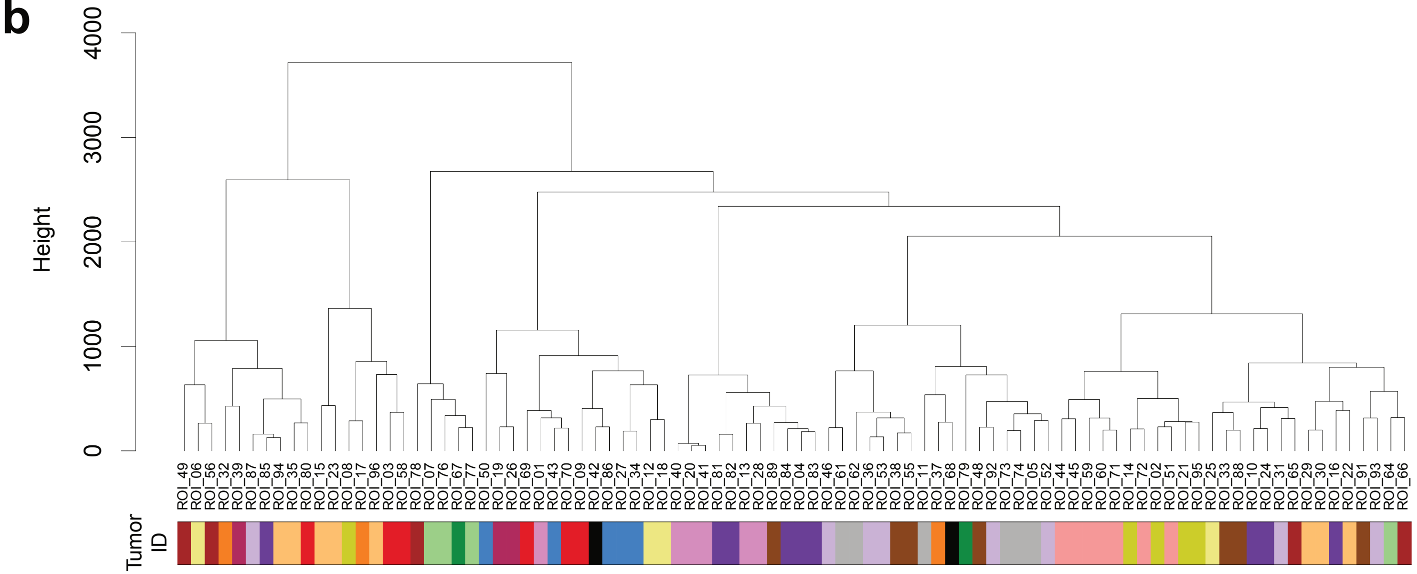

#### Supplemental Fig. 2

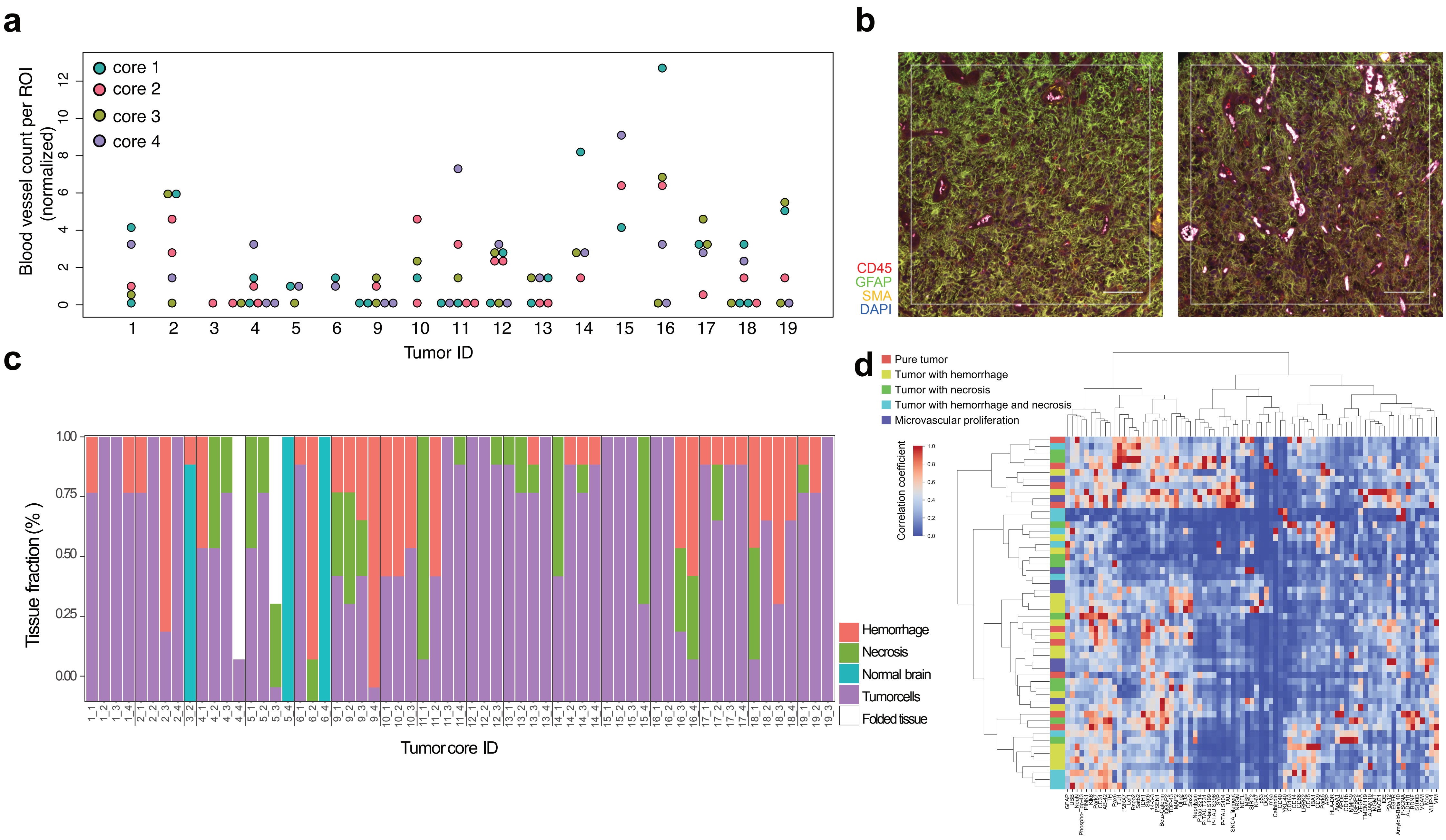

Supplemental Fig. 3

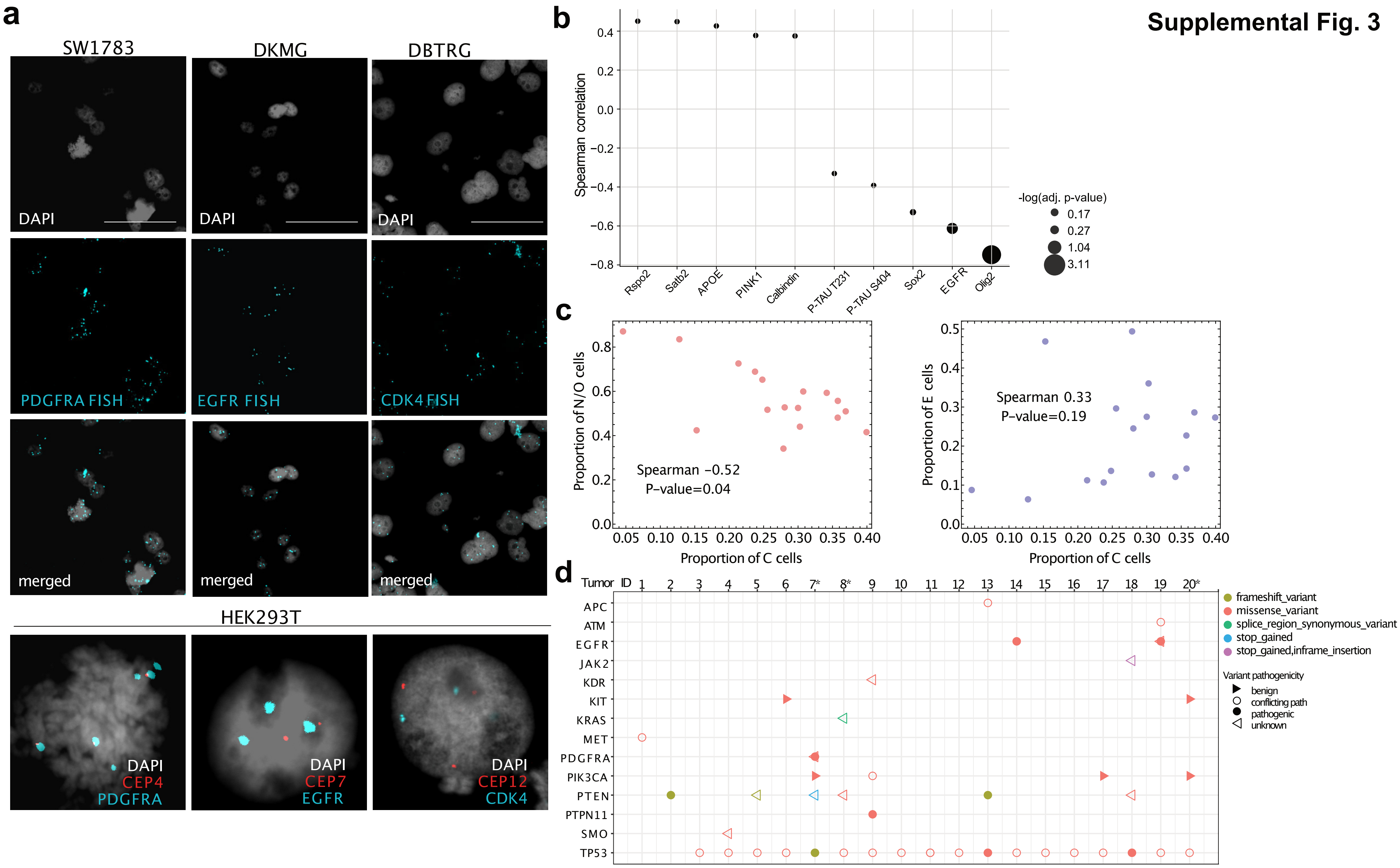

#### Supplemental Fig. 4

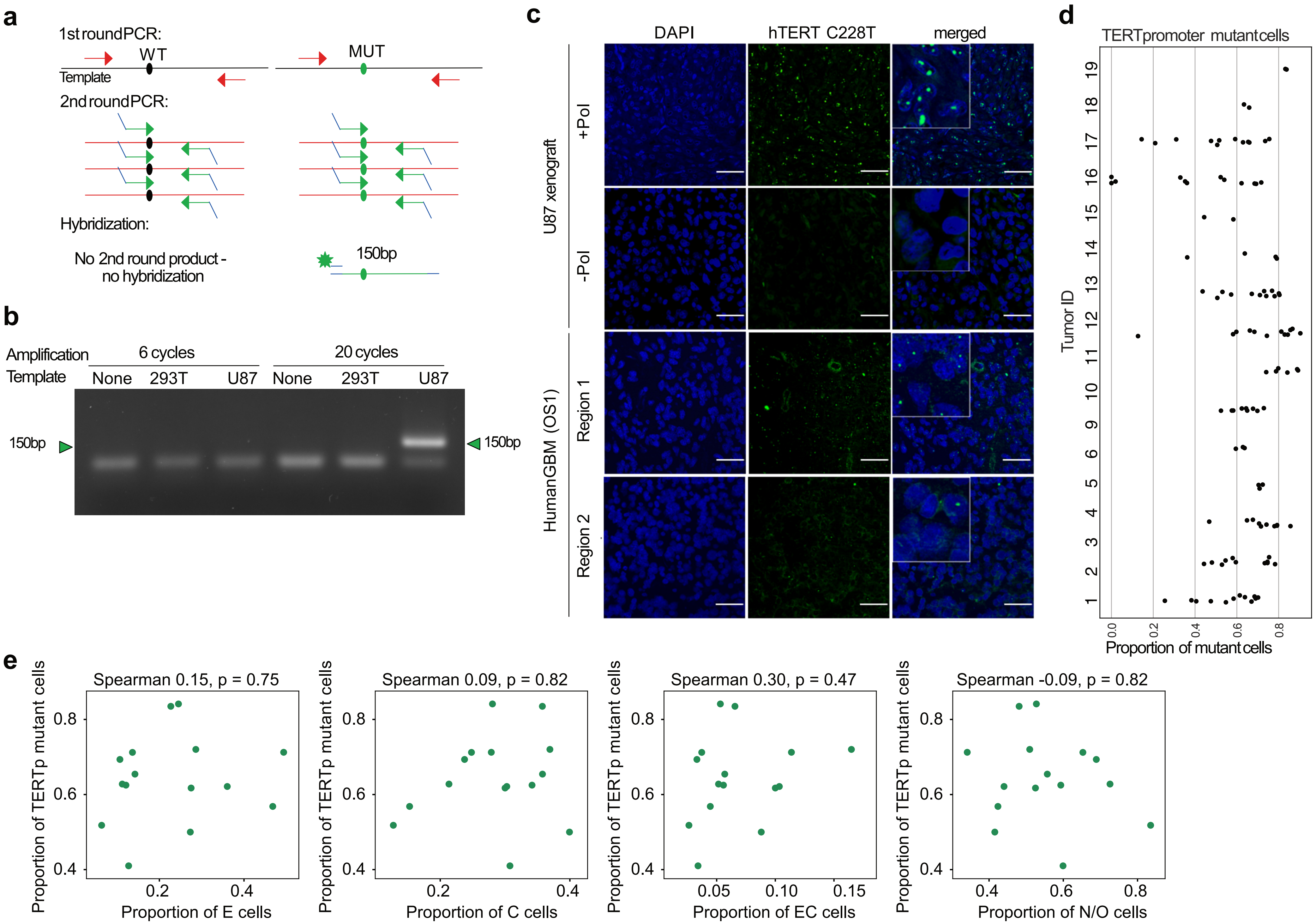
